# Fast assembly and *in vivo* coalescence of ParB_F_ biocondensates involved in bacterial DNA partition

**DOI:** 10.1101/2025.10.27.684735

**Authors:** Perrine Revoil, Linda Delimi, Jérôme Rech, Josh Cailhau, François Cornet, Jean-Charles Walter, Jean-Yves Bouet

**Affiliations:** Laboratoire de Microbiologie et Génétique Moléculaires, Centre de Biologie Intégrative (CBI), Centre National de la Recherche Scientifique (CNRS), Université de Toulouse, F-31062 Toulouse, France; Laboratoire Charles Coulomb (L2C), CNRS, Université de Montpellier, Montpellier, France

**Keywords:** ParABS system, Partition complex, ParB, DNA segregation, Biomolecular condensates, Condensate coalescence

## Abstract

Faithful DNA segregation in bacteria relies on ParABS systems, in which ParB assembles into condensates at centromere-like *parS* sites, while the ATPase ParA spatially organizes these complexes. How these ParB condensates maintain dynamic behavior without collapsing into a single structure has remained unclear. Here, we combine chromosome degradation with quantitative imaging to dissect the kinetics and physical principles governing ParB condensate dynamics *in vivo*. In the absence of the nucleoid, ParB condensates from the plasmid F diffuse freely and coalesce within seconds upon encounter. Strikingly, quantitative analyses indicate that condensates may operate near the fusion-separation boundary, such that minimal energy is sufficient to split them after replication, preventing irreversible coalescence. Using mutants, we demonstrate that condensate assembly is required for coalescence and uncover a dual role for ParA_F_: nucleoid tethering restricts condensate mobility and limits fusion, while ParA_F_ also promotes a ParB_F_ state competent for assembly and coalescence, likely by enhancing ParB-ParB interactions. Finally, condensates rapidly disassemble and reassemble upon 1,6-hexanediol treatment, underscoring their reversibility and the stabilizing contribution of ParB-DNA interactions. Together, our results establish ParB_F_ complexes as *bona-fide* biocondensates tuned by ParA_F_ to ensure robust DNA segregation. More broadly, these findings highlight regulated phase separation as a key organizing principle of bacterial replicons.

## Introduction

Bio-molecular condensates, also referred as membrane-less organelles, are key organizational units in living systems, facilitating a wide range of cellular processes (1,2). They have been widely reported in eukaryotes, and more recently in bacteria where they concentrate proteins and nucleic acids without the need for a lipid membrane (3,4). In bacteria, such condensates contribute to diverse processes, including transcription (5), mRNA decay (6), and the partitioning of intracellular cargos, such as carboxysomes in autotrophic bacteria (7,8) and DNA centromeres from plasmids and chromosomes (9–11).

Active DNA partitioning in bacteria is primarily driven by the ParABS system, widely found on low-copy-number plasmids and the only one present on chromosomes (12). This system is composed of three components: two proteins, ParA and ParB, and a centromeric sequence, *parS*. Together, these elements are necessary and sufficient to ensure accurate plasmid inheritance. ParB assembles into higher-order nucleoprotein complexes at *parS* sites, while ParA, an ATPase, spatially localizes these partition complexes through dynamic interaction with the nucleoid. In its ATP-bound form, ParA associates with DNA and interacts with the cargo via ParB, which stimulates its ATPase activity and promotes its release from DNA (13–17). This interaction generates local ParA depletion zones on the nucleoid, driving directional cargo movement toward regions of higher ParA concentration through a diffusion-ratchet mechanism (18–21). In the plasmid F system, a subset of ParA_F_ also associates with high-density-regions of the nucleoid and contributes to anchoring ParB_F_ condensates within the nucleoid (21,22). Following DNA replication and duplication of ParB-*parS* complexes, ParA mediates their separation and relocation to opposite cell halves, ensuring faithful inheritance upon cell division (12,23).

Partition complexes regroup over 80% of ParB at centromere sites (14,24). Their assembly are initiated by a sequential multi-step process: (i) specific binding of ParB to *parS*, typically a 16-bp DNA motif (25–27), (ii) CTP binding to *parS*-bound ParB (28–30), (iii) conversion of ParB into a sliding clamp, promoting its release from *parS* (30–32), (iv) diffusion at short distances along *parS*-proximal DNA (30,33), and (v) clamp opening followed by ParB unloading (29,34). ParB thus cycles between open and clamped forms, called the ParB-CTP cycle, enabling the loading of ParB clamps specifically at *parS* sites. Notably, both chromosomal and plasmid ParABS systems form partition complexes that display hallmarks of biomolecular condensates (9,11), characterized by a high-density ParB phase coexisting with a dilute phase, likely arising from phase separation (4,35). This behavior has been proposed to rely on a CTP-dependent molecular switch that enhances interactions between ParB dimers (36–38), thereby driving strong local enrichment of ParB (9,10,24,39). In addition, ParB clamp-mediated bridging between distant DNA segments may further stabilize these assemblies (10,24,39–42), forming the higher-order nucleoprotein complexes that dynamically interact with ParA to ensure the segregation of chromosomal origins and plasmids (23).

Biomolecular condensates typically exhibit liquid-like properties, including spherical shapes, rapid exchange of components between condensed and diffuse phases, and coalescence into larger droplets (43). Partition complexes in the plasmid F system displays these features *in vivo*, including rapid exchange of ParB_F_ between complexes (44), reduced intracellular mobility of ParB_F_ within rounded condensates (∼100-fold lower than outside) (9), and fusion events upon ParA_F_ depletion (9). At thermodynamic equilibrium, in a finite and confined system, phase separation with a condensed phase minimizes interfacial free energy by forming a single condensate, with coalescence or Ostwald ripening driving this process (45). However, in ParABS systems, this intrinsic tendency toward coalescence conflicts with the functional requirement to maintain multiple, spatially separated ParB condensates to ensure faithful DNA partition. Although ParA_F_ depletion induces condensate fusion (9), these events occur slowly, likely due to sustained high expression of ParA_F_ prior to its gradual degradation, limiting quantitative analysis of coalescence dynamics.

Here, we developed an inducible chromosome degradation system to probe the behavior of ParB condensates in the absence of the nucleoid. Under these conditions, ParB_F_ condensates diffuse freely and coalesce within seconds upon encounter, enabling direct measurement of coalescence kinetics. We further show that condensate assembly is rapid, reversible, and dependent on hydrophobic interactions, as revealed by hexanediol sensitivity. Importantly, our results uncover a previously unappreciated function of ParA_F_, beyond nucleoid tethering, in promoting ParB_F_ condensate assembly. Together, these functions regulate the fusion-fission balance of ParB_F_ condensates, maintaining their dynamic stability and ensuring robust and faithful segregation of bacterial DNA.

## Material and methods

### Bacterial strains, plasmids and oligonucleotides

All strains used in this study are derivatives of *E. coli* K12 and are listed, together with plasmids in Table S1. Details of plasmid and strain construction are provided in the Supplementary experimental procedures. Briefly, strain LY643, which harbors four I-SceI sites (gift from C. Lesterlin), was modified to allow uniform induction of *para_BAD_::I-sceI* by introducing the *Pcp18::araE* alleles (46) via P1 transduction. To enhance chromosome DNA degradation efficiency, a *recA* mutation was also introduced, resulting in strain DLT4151.

All F and mini-F plasmids are derivatives of F1-10B (44) and pDAG114 (46), respectively. F1-10B derivatives were introduced into recipient strains by conjugation, while pDAG114 derivatives were introduced by CaCl_2_**-**transformation. The plasmid producing I-SceI, pSN01 (47), was introduced last by CaCl_2_**-**transformation.

Oligonucleotides are listed in Table S2.

### Epifluorescence microscopy

Strains expressing fluorescently tagged proteins were grown overnight at 30°C in M9 minimal medium supplemented with glucose and casamino acids (M9-glucose-CSA). Cultures were diluted 1:250 in fresh medium and incubated at 30°C until OD_600nm_ ∼0.3. Cells were then either spotted (0.7 μl) onto M9-buffered 1% agarose pads or loaded into microfluidic devices (Chitozen slides, Idylle). Where indicated, 0.2% arabinose was added to cell cultures to induce I-SceI expression. Elongated *E. coli* cells were obtained by adding cephalexin (10 µg.ml^-1^) to the growing cultures 30 min. prior to arabinose induction and further microscopy observations.

Fluorescence images were acquired as previously described (48) using a Nikon epifluorescence microscope and NIS-Elements AR software (Nikon) for image capture and processing. Image analysis was performed using ImageJ softwares, with foci detection and positioning on the longitudinal cell axis carried out using the “*Coli inspector*” macro and the “*ObjectJ*” plugin, respectively.

For nucleoid staining, cells were incubated with DAPI (1 µg.ml^-1^) for 10 min. under shaking prior to loading into microfluidic slides, followed by continuous flow with M9-glucose medium containing DAPI (1 µg.ml^-1^).

To quantify the ratio of ParB fluorescence intensity before and after fusion events, cells carrying two visible ParB condensates were selected. Regions of interest (ROIs) were manually drawn around each condensate in three consecutive frames before fusion, and the total fluorescence intensity was averaged. After fusion, a single ROI was drawn around the merged condensate and the fluorescence was averaged over three consecutive frames. Background fluorescence from the medium was subtracted from all intensity measurements.

### Skewness analyses

Individual cells were identified on phase-contrast images by cell segmentation using MicrobeJ. The resulting masks were subsequently applied to the corresponding fluorescence images for quantitative analysis. Skewness measurements, calculated using MicrobeJ plugin “*skewness*”, measures the asymmetry of the pixel intensity distribution relative to the average fluorescence within each cell, providing a quantitative readout of condensate presence and compactness. Skewness values for each cell were automatically extracted and plotted using the BoxPlot function in the MicrobeJ plugin, based on the intensity.channel.skewness parameter.

### Mean square displacement

Exponential growing cells, induced or not with 0.2% arabinose, were spotted onto 1% agarose pads buffered with M9-glucose medium. Time-lapse fluorescence microscopy was performed with images every 500 milliseconds over a 30-second period. ParB-mVenus foci were detected and tracked using the *TrackMate* plugin (49) in Fiji. For each track, the position of the foci was recorded over time, and the MSD was calculated as a function of time lag. The diffusion coefficient (D) was estimated from the slope of the initial linear portion of the MSD curve, assuming Brownian diffusion. All fitting and statistical analyses were performed using custom scripts (See Supplementary material and methods for details).

### Physical modeling

Details of the theoretical physics employed for physical modeling are described in Supplementary data. It includes the description of polymer model and coarse-graining analyses, the calculation of the Critical surface tension (γ_crit_) and of the free-energy equilibrium.

## Results

### Chromosome depletion triggers coalescence of ParB condensates

ParB_F_ condensates are typically anchored within the nucleoid via interactions with ParA_F_, which binds to the high-density regions of DNA (22). To investigate the dynamics and conditions that promote fusion of ParB_F_ condensates, we developed a strategy to disrupt this anchoring by selectively removing the bacterial chromosome. We hypothesized that depleting the nucleoid would free ParB_F_ condensates to diffuse throughout the cytoplasm, facilitating their coalescence, if any. To test this, we employed a method based on the reckless phenotype for specific and progressive chromosome degradation while preserving plasmid integrity (47,50). In the absence of RecA, which is essential for homologous recombination, induction of double-strand breaks (DSBs) leads to complete chromosome degradation by the RecBCD helicase-nuclease complex. We constructed an *E. coli recA* strain (DLT4151) carrying four I-SceI restriction sites evenly distributed across the chromosome (Fig. 1A). This strain was transformed with pSN01, which expresses the I-SceI endonuclease under the control of arabinose, enabling inducible DSBs and subsequent chromosome degradation. The amount of nucleoid per cell was monitored over time using DAPI staining and fluorescence microscopy (Fig. 1B, left panels). Upon arabinose-induced I-SceI production, DAPI fluorescence intensity gradually decreased to near-baseline levels over ∼100 minutes, confirming progressive loss of chromosomal DNA (Supplementary Fig. S1A-C). By 120 minutes post-induction (T120), chromosomal DNA was extensively degraded in nearly all cells.

**Figure 1:**
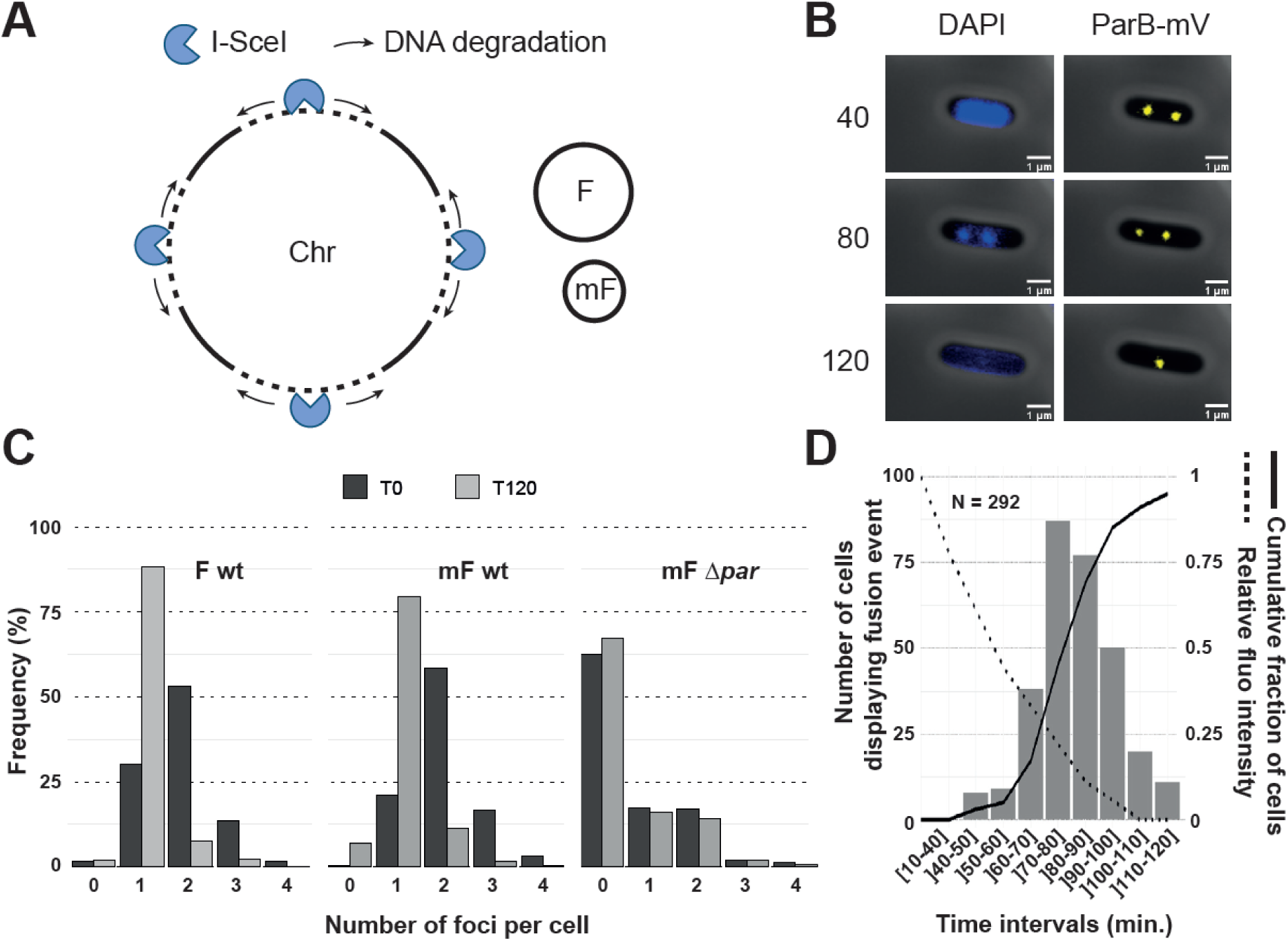
ParB partition condensates coalescence in the absence of nucleoid. **A-** Schematic representation of the inducible chromosome degradation system. I-Sce1 endonuclease cuts at the four specific sites. RecBCD complexes specifically degrade the chromosome but not plasmid DNA from the double strands breaks (black arrows). **B-** Fluorescence microcopy of chromosome degradation and fusion events. Series of images of the same cell, stained with DAPI, at 40, 80 and 120 min. post-arabinose induction, observed in the blue (left panels) and yellow (right panels) channels, to monitor chromosome degradation and ParB-mVenus condensates. Scale bar, 1 µm. **C-** Distribution of the number of ParB-mVenus foci. The number of foci per cell before (T0; black) and after (T120; grey) chromosome degradation is displayed for the plasmid F (F1-10B04; left), the miniF (pJYB234; middle) and the miniF Δ*parAB* (pPER07; right). **D-** Distribution of ParB-mVenus fusion events over time. Images (as in B) were acquired every two minutes between 40 and 120 min. after arabinose induction. The number of cells displaying a fusion event were shown per time intervals. The solid line represents the cumulative fraction of chromosome-less cells displaying a fusion event. The dotted line represents the degradation of the chromosome over time (from Supplementary Fig. S1B). N corresponds to the number of cell analyzed.

We then examined the behavior of ParB_F_ condensates in the absence of the nucleoid. F plasmids producing ParB_F_-mVenus fusion proteins were introduced in the DLT4151 strain, which was further transformed with pSN01. In steady-state growth, prior to induction (T0), cells carrying the 100-kbp F plasmid (F1-10B04) typically exhibited two ParB_F_-mVenus foci. After chromosome degradation (T120), 88.4% of cells contained only a single focus (Fig. 1B, right panel and Fig. 1C, left panel). To directly observe these fusion events, we performed time-lapse imaging at 2-minute intervals. Fusion was defined as the merging of two foci that subsequently moved together during several successive images (Fig. 1D). Most fusion events occurred between 70 to 90 minutes post-induction, corresponding to ∼75-90 % of chromosome degradation (dotted line). By T120, when chromosome degradation was complete, fusion events had occurred in 95% of the cells, as shown by the cumulative curve (black line).

To assess whether plasmid size affects condensates fusion, we repeated the experiment using a ∼10-kbp mini-F plasmid (pJYB234). Similar coalescence kinetics were observed, with over 90 % of the cells exhibiting fusion events, as with the 100-kbp plasmid (Fig. 1C, central panel and Supplementary Fig. S1D). This finding indicates that condensate fusion is independent of plasmid size.

To ensure that plasmids do not spontaneously cluster in the absence of the chromosome, we monitored the behavior of a mini-F lacking the *par* system, labelled with a FROS system allowing us to track plasmids devoid of partition complexes (Fig. 1C, right panel). Due to defective partitioning, this Δ*par* mini-F (pPER07) was frequently lost, resulting in a high proportion of plasmid-free cells, *i.e.* cells without foci (62%). Among cells retaining the plasmid, the proportions with one (17.3%) or two (16.9%) foci remained unchanged before (T0) and after complete chromosome degradation (T120). These results demonstrate that, in nucleoid-free cells, low-copy-number plasmids do not cluster together in the absence of the ParABS system.

### Defects in ParB multimerization impair condensate fusion

To determine whether the fusion events observed in nucleoid-free cells reflect an intrinsic property of ParB_F_ condensates, we analyzed a ParB variant, ParB_F_-R156A, previously shown to partially impair condensate formation and partition activity (36). This mutation, located in the box III motif involved in CTP binding and hydrolysis (31) still allows ParB_F_-R156A to bind to *parS*_F_ and to be released as sliding clamps, but it is unable to fully assembled the higher-order assembly of ParB_F_, as evidenced by ChIP-sequencing analyses of the ParB_F_-R156A DNA binding profile (36). We monitored cells carrying an F plasmid expressing ParB_F_-R156A-mVenus before (T0) and after I-SceI induction (Fig. 2A, top panel). At T0, 15% of the cells lacked detectable foci, consistent with plasmid loss due to defective partitioning (Fig. 2A, bottom panel), compared to 1.5 % in the wild-type (Fig. 1C). At T120, a shift from two foci to one was observed, indicating that coalescence can occur. However, this effect remained limited: the proportion of one-focus cells increased by ∼18%, and two-focus cells decreased by ∼19%, in contrast to 58% and 47%, respectively, for the WT. Moreover, at T120, only 50 % of chromosome-depleted cells exhibited fusion events, and with delayed kinetics (Supplementary Fig. S1E), compared to ∼95% in wild-type (Fig. 1D). These results indicate that partial defect in ParB_F_ condensate assembly substantially reduces the frequency and efficiency of their fusion. Consistent with this defect, ParB_F_-R156A-mVenus displayed a markedly diffuse fluorescence pattern with less-defined condensates compared to the ParB_F_-mVenus (Fig. 2A versus Fig. 1B). To quantify this difference, we performed a skewness analysis of fluorescence intensity distributions, which reports on the asymmetry of intracellular signal. Higher skewness values correspond to a few bright pixels for a majority of low intensity pixels, or punctate foci (*i.e.*, well-formed condensates), whereas close to zero values indicate diffuse fluorescence. This metric thus provides a quantitative proxy for ParB condensates assembly (Fig. 2B). In WT plasmid, ParB_F_-mVenus formed bright foci with an average skewness of 1.85 (N=144). In contrast, cells carrying a mini-F Δ*parS* plasmid, which is unable to nucleate the formation of ParB condensates, displayed near-baseline skewness (0.19; N=411). The *parB*_F_-R156A condition showed intermediate skewness values (0.70; N=269), indicating defective condensate assembly. These results support the conclusion that impaired ParB self-association compromises both condensate formation and their ability to undergo fusion.

**Figure 2:**
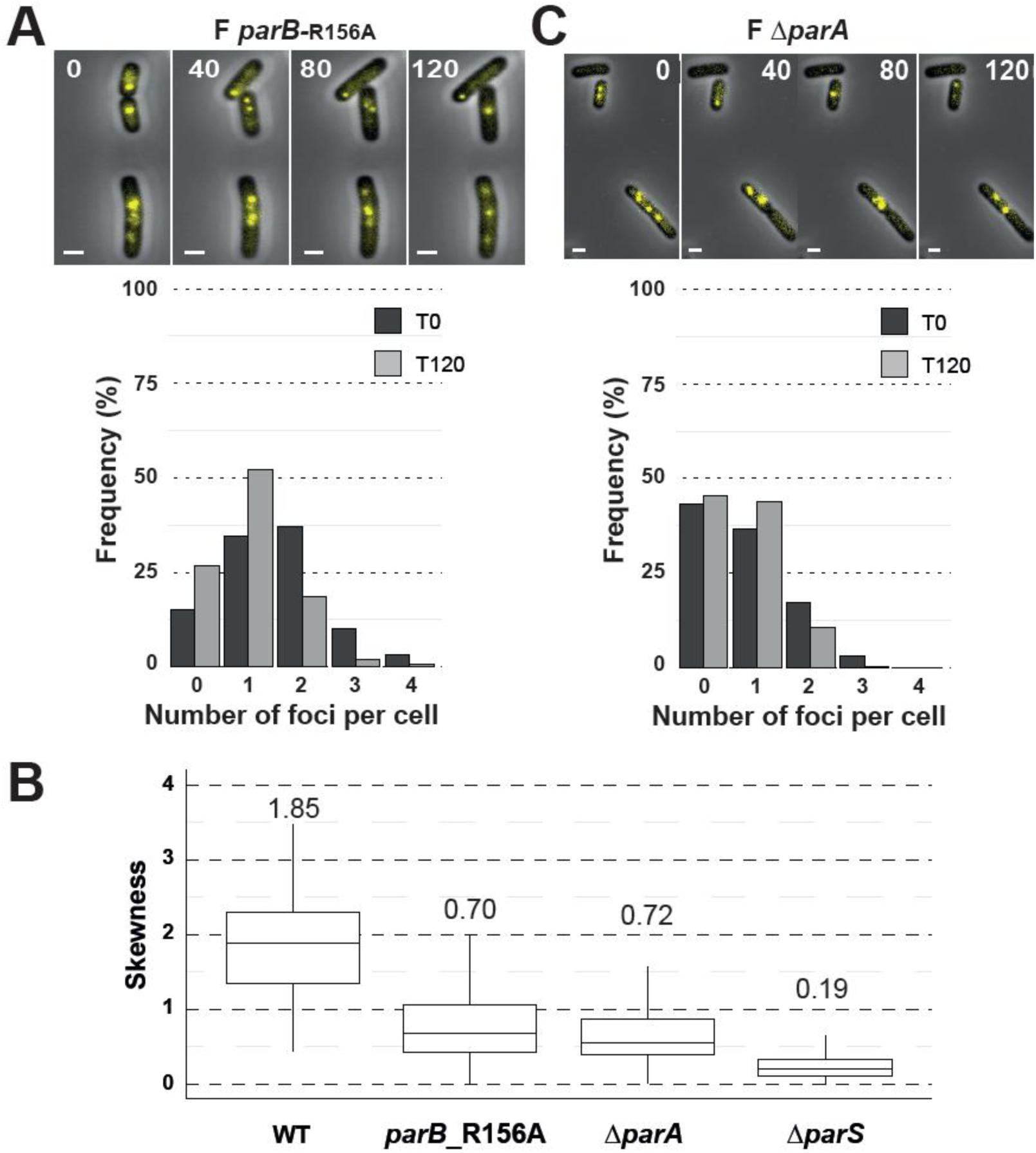
Coalescence depends on properly assembled ParB condensates. **A**- Time-lapse imaging of ParB_F_-R156A-mVenus. (top) Series of images acquired at 0, 40, 80 and 120 minutes post-arabinose induction (plasmid F1-10B15). Scale bar, 1 µm. (bottom) Distribution of the number of ParB-R156A-mVenus foci before (T0; black) and after (T120; grey) chromosome degradation. **B**- Time-lapse imaging of ParB_F_-mVenus from the F Δ*parA* plasmid. As in (A), with the plasmid F1-10B12. **C**- Quantification of the ParB condensates assembly defect. Fluorescent images of ParB_F_-mVenus or ParB_F_-R156A-mVenus, from WT (N=144), Δ*parA* (N=152), Δ*parS* (N=411) or *parB*-R156A (N=269) plasmids in exponentially growing cells (no chromosome degradation) were subjected to Skewness analyses. Skewness analyses were displayed as box plots indicating the medians, the quartiles and the standard deviations, and average values are indicated on top of each box plot.

### ParA_F_ has an effect on the coalescence of ParB_F_ condensates

ParA_F_ plays a central role in positioning partition complexes along the nucleoid (21,22). In its ATP-bound form, ParA_F_ interacts with both ParB_F_ and non-specific DNA (nsDNA) (51,52), and has been proposed to promote ParB_F_-ParB_F_ interactions (42). To test whether ParA_F_ also contributes to condensate coalescence, we analyzed a plasmid F Δ*parA*_F_ (F1-10B12), expressing ParB_F_-mVenus from an inducible P*tet* promoter. Western blot analysis indicated that ParB_F_-mVenus levels were about two-fold lower than in the wild-type, where expression is driven by the native ParA_F_ auto-regulated promoter (Supplementary Fig. S2). Despite this reduction, partition condensates were readily detectable (Fig. 2C, top). We quantified the number of ParB foci per cell before (T0) and after (T120) I-SceI induction (Fig. 2C, bottom). At T0, 43% of the cells lacked foci, consistent with defective partitioning in the absence of ParA_F_ (22). Following chromosome degradation (T120), only a modest shift was observed: the proportion of one-focus cells increased by 7%, while two-focus cells decreased by 6.4%. This contrasts with the wild-type condition (45% decrease in 2-foci cells; Fig. 1D), indicating that the absence of ParA_F_ strongly impairs coalescence of ParB condensates in the absence of chromosome.

ParB fluorescence in cells carrying a Δ*parA*_F_ plasmid appeared more diffuse (Fig. 2C), similar to that observed for the ParB_F_-R156A variant (Fig. 2A). Skewness analysis yielded an intermediate value (0.72; N=152) (Fig. 2B), comparable to that obtained with ParB_F_-R156A. This partial defect likely reflects impaired ParB_F_-ParB_F_ interactions upon loading at *parS*_F_, preventing efficient condensation of the majority of ParB_F_.

Together, analyses of the *parB*_F_-R156A mutant and the Δ*parA*_F_ condition indicate that, in the absence of the nucleoid, condensate coalescence is severely impaired when ParB self-association is disrupted. In both cases, reduced fusion correlates with defective condensate assembly, indicating that efficient coalescence requires a properly assembled condensate state. We therefore propose that ParA_F_, in addition to its role in spatial organization, may promote ParB condensate assembly, thereby enabling their fusion in the absence of chromosome.

### Coalescence into a single condensate is a conservative process

Condensates naturally tend to fuse into fewer, larger assemblies, as droplet coalescence minimizes surface tension and free energy, thereby reaching a more thermodynamically stable state (45). However, the diffraction limit of fluorescence microscopy prevented us from directly measuring the size of individual ParB condensates before and after fusion. To overcome this technical limitation, we used fluorescence intensity as a proxy for condensate size. This approach relies on our previous finding that the ParB concentration inside condensates is comparable to the overall protein concentration in the cytoplasm, suggesting that condensates accumulate ParB up to a cytoplasmic saturation threshold (9). Under these conditions, fluorescence intensity is thus expected to scale with condensate volume.

We measured the fluorescence intensity of ParB-mVenus foci in 50 cells, before and after fusion events. On average, the post-fusion intensity was 0.94 times the sum of the two pre-fusion foci (Fig. 3A and Supplementary Fig. S3). This indicates that the majority of ParB molecules are retained within the resulting condensate, consistent with a conservative coalescence process in which little to no ParB is lost or dispersed. This finding supports the idea that fusion results in the formation of larger condensates.

**Figure 3:**
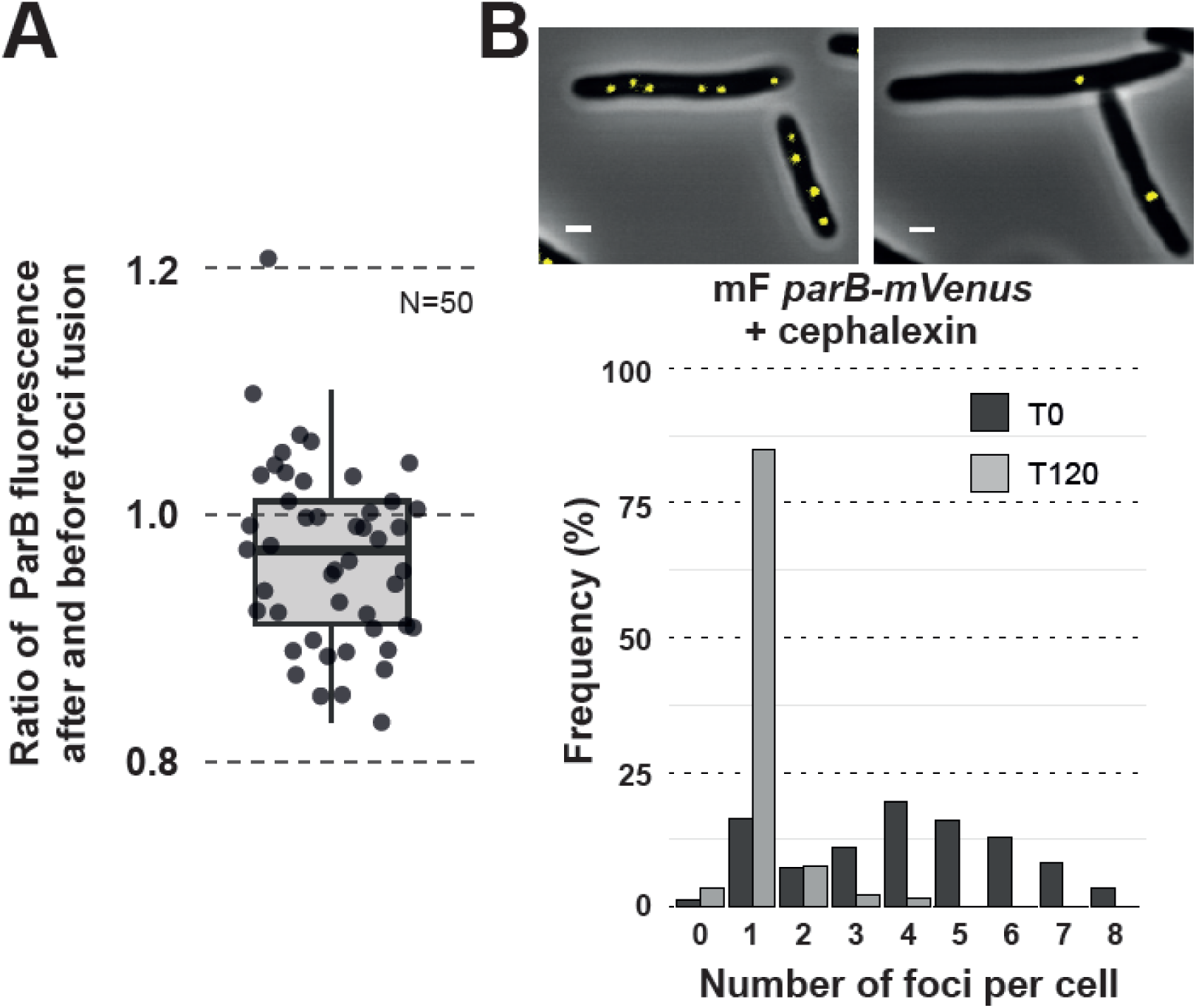
Coalescence into a single condensate is conservative. **A**- Most ParB molecules regroups in the fused condensates. The box plot displays the ratio of the average ParB fluorescence intensity, measured on three successive images, after fusion events over the average sum of ParB-mVenus fluorescence intensity from two foci, measured on three successive images, before fusion events. The box plots indicates the median, quartiles and the standard deviation, with each dot representing a fusion event (N=50). **B**- Numerous ParB condensates coalesce into a single one after chromosome degradation. (top) Imaging of ParB_F_-mVenus from strain DLT 4161 (pJYB234), grown in the presence of cephalexin, after 40 and 120 min post-arabinose induction. The scale bar represents 1 µm. (bottom) Distribution of the number of ParB-mVenus foci before and after chromosome degradation. The number of foci per cell before (T0; black) and after (T120; grey) chromosome degradation is displayed for the plasmid mF pJYB234.

We next asked whether more than two foci could coalesce into a single focus. To test this, we treated cells with cephalexin, a β-lactam antibiotic that blocks cell division without inhibiting DNA replication. This treatment induced cell elongation and increased the plasmid copy number per cell. After one hour of cephalexin exposure, cells had approximately doubled in size, and ParB foci ranged from 1 to 8, with an average of 4.2 foci per cell (Fig. 3B). Following chromosome degradation, 88% of cells exhibited a single ParB focus at T120, indicating that most ParB condensates had coalesced into a single one.

These findings support the view that ParB condensates behave as droplets undergoing coalescence. Regardless of plasmid copy number, ParB condensates reliably fuse into fewer, larger assemblies to minimize surface tension. This process is largely conservative, retaining all ParB molecules within the fused condensates.

### ParB-DNA condensates operate near the fusion-separation boundary

Our measurements showed that 95 % of chromosome-depleted cells had undergone partition condensates fusion within 120 min. of chromosome degradation induction (Fig. 1E). We were curious about the remaining ∼5% of cells in which fusion was not observed at T120. Closer inspection revealed that some cells displaying a single fused focus at earlier time points latter exhibited splitting events. To explore this phenomenon, we performed high-frequency imaging, acquiring frames every 250 milliseconds over 30-second (Fig. 4A). Despite this brief time window, we detected splitting events in ∼10 % of cells carrying the plasmid F (Fig. 4B), suggesting that splitting is a frequent event, potentially occurring approximately once every five minutes per cell on average. When examining cells carrying the smaller mini-F, splitting events were detected in 18% of cells, indicating that condensates derived from smaller plasmids split more frequently than those from larger plasmids. This suggests that, in the absence of chromosome, a smaller DNA amount near *parS* sites permits more frequent escape of one plasmid from condensates. In both cases, splitting events were transient, as separated foci consistently fused back after a short period of time (Fig. 4A). Notably, the slightly higher splitting frequency observed in Δ*parA* derivatives, compared to their respective wild-type counterpart (Fig. 4B), was not statistically significant according to our tests.

**Figure 4:**
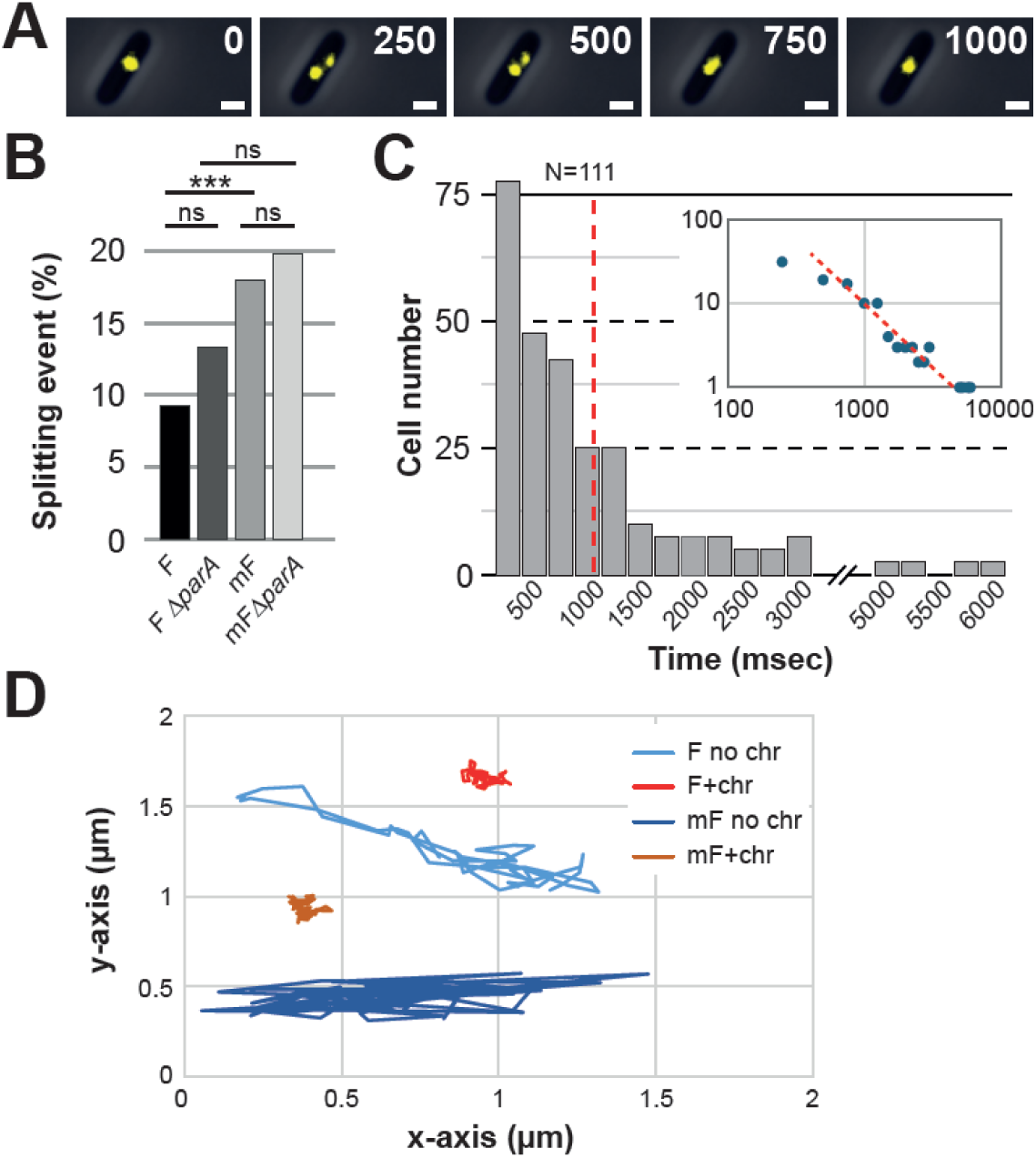
Coalescence of partition condensates is nearly instantaneous. **A**- Transient splitting of ParB condensates in chromosome-depleted cells. Time-lapse fluorescence microscopy was performed after complete chromosome degradation (time > T120), with images acquired every 250 msec. Time is indicated in msec. Transient splitting and re-fusion of ParB-mVenus foci were frequently observed. **B**- Splitting events are frequent. The percentage of splitting events was quantified in chromosome-depleted cells expressing ParB-mVenus from plasmid F or mini-F (mF), in the presence or absence of *parA* (Δ*parA*). N= 807 (F), 283 (mF), 120 (FΔ*parA*) and 86 (mFΔ*parA*). Pairwise comparisons were performed using Fisher’s exact test and Chi^2^ test. P-values were adjusted for multiple comparisons using the Bonferroni correction (α = 0.0125 for 4 comparisons). Results were consistent using both Chi² and Fisher’s exact tests (Bonferroni-corrected). ns (not significant, P ≥ 0.0125); *** (P < 0.001). **C**- Fusion occurs rapidly following splitting. A total of 111 chromosome-depleted cells carrying plasmid F and exhibiting a splitting event (as in A) were analyzed. The time elapsed between splitting and fusion was measured, revealing that fusion typically occurred within ∼1 second. (*inset*) same data plotted in Log-Log scale (blue dots). The dotted red line corresponds to a power law with an exponent -3/2. **D-** Partition condensates are highly mobile in the absence of the chromosome. Representative intracellular trajectories of ParB-mVenus foci tracked over time in cells carrying plasmid F or mini-F, either with (+chr) or without (no chr) the chromosome. Trajectories illustrate an important increase in mobility under chromosome-depleted conditions.

In nucleoid-depleted cells, condensates are fused ∼90% of the time but still undergo occasional splitting, suggesting that the system operates near a crossover regime between fused and separated states, with a free energy difference approaching zero, *i.e.*, within the fused region of the phase diagram but close to the transition line (Δ*F* ∼ 0). To quantitatively understand this equilibrium, we developed a coarse-grained polymer model that captures the competition between surface energy minimization and configurational entropy loss during condensate coalescence (see Supplementary Methods). At the fusion-separation transition, we estimated a critical surface tension γ_crit_ = 0.80 +/- 0.21 µN.m^-1^ for the F plasmid (*N* = 660; Supplementary Fig. S4). Given that ParABS systems are naturally found on replicons spanning a wide range of size, from ∼25 kbp to over 1 Mb (53), we extended our analysis and calculated a γ_crit_ = 1.10 +/- 0.29 µN.m^-1^ for a 1 Mb DNA molecule. For the smaller mini-F used in this study (pJYB234, ∼10 kbp; *N* = 60), we measured a γ_crit_ = 0.49 +/- 0.13 µN.m^-1^ (Supplementary Fig. S4). These values place the tension surface of ParB-DNA droplets, across a large range of DNA size replicons, within the typical 0.1-1 μN.m^-1^ range observed in biological phase-separated systems (54). According to both the experiments and the model, which captures the fundamental symmetries of the system, we hypothesize that ParB-DNA condensates operate near the fusion-separation boundary, with entropy costs remaining negligible over biologically relevant DNA lengths. Further studies will be required to assess the impact of neglected effects such as plasmid supercoiling.

### Coalescence of partition condensates occurs rapidly

The dynamic splitting-fusion behavior observed for ParB foci provided an opportunity to investigate the kinetics of condensate coalescence. Using the frame where a splitting event was first observed as time zero, we measured the time required for ParB condensates to fuse back together. Analysis of 111 such events from the F plasmid revealed that re-fusions occurred, on average, within approximately one second (Fig. 4C). This result demonstrates that coalescence of partition condensates is a rapid process, nearly instantaneous compared to the time of splitting, the cell cycle or to DNA replication. To further characterize the fusion kinetics, we plotted the number of fusion events over time on a log-log scale and found that the distribution fit well to a power law with an exponent of -3/2 (Fig. 4C, inset). This scaling suggests that the coalescence times are consistent with a first encounter of two Brownian particles (55). Such behavior is consistent with free diffusion of condensates in the absence of chromosome, where condensate collisions and subsequent fusions are governed by random Brownian motion.

To assess the mobility of partition condensates, we compared their diffusive behavior in the presence and absence of the chromosome. ParB_F_-mVenus foci from both plasmid F and mini-F were tracked using high-resolution time-lapse microscopy, with images acquired every 0.5 second over a 30 seconds period. In chromosome-depleted cells, condensates exhibited markedly enhanced mobility (Supplementary Fig. S5): the mean step size (*Ss*) between frames increased by 6-fold for mini-F (*Ss* ∼ 1.8 µm) and 4-fold for plasmid F (*Ss* ∼ 1.2 µm) compared to cells with intact chromosomes (*Ss* ∼ 0.3 µm). We quantified this difference by computing the mean square displacement (MSD) of ParB_F_ foci to estimate diffusion coefficients (*D*) under both conditions (Fig. 4D and Supplementary Fig. S5), using a sub diffusive model:

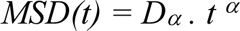

where D*α* is the generalized diffusion coefficient (µm² s^−^*^α^*) and *α* is the scaling exponent describing the nature of particle motion (see Eq. (5) in Supplementary data). In the presence of the chromosome, α values of 0.81 (F) and 0.88 (mini-F) reflected sub-diffusive, constrained dynamics (56). Upon chromosome removal, *α* rose to 0.92 and 0.95, respectively, approaching the theoretical value for Brownian (free) diffusion (*α* = 1). The corresponding D*_α_* values increased by 10-40 fold, from 0.001-0.002 µm² s^-α^ to 0.04 and ∼0.02 µm² s^-α^. Notably, the rise in *D_α_* scales approximately with the square of the step size, confirming that the exponents *α* are sufficiently close to 1 to approximate *D_α_* to an usual diffusion coefficient at short time scales. Together, these results show that ParB_F_ condensates behave as nearly freely diffusing particles in the absence of chromosome, and suggest that their mobility is primarily limited by condensate size rather than the length of the plasmid DNA carrying *parS*_F_.

### Partition condensates assemble and disassemble within seconds

In biological phase separation, biomolecular interactions, such as protein-protein and protein-nucleic acid interactions, underpin the formation and stability of condensates. The ParB condensates formed on plasmid F enables the dissection of the individual roles of these two interactions within cells almost completely depleted of non-specific DNA. To test the role of weak intermolecular interactions in condensate stability, we used 1,6-hexanediol (Hex), an aliphatic alcohol that interferes with hydrophobic interactions (57).

Cells expressing ParB_F_-mTq2 from mini-F along with a FROS reporter MalI-mNG^2^ (bound to 7 *malO* sites present on the plasmid; pPER08) were deprived from their chromosome and, when ∼95% of cells displayed fused partition condensates (T120), were treated with 10 % Hex. After one minute, ParB-mTq2 fluorescence became diffuse in most cells (89.3%), indicating efficient condensates dissolution (Fig. 5A and Supplementary Fig. S6A). In contrast, MalI-mNG^2^ foci remained intact, showing that specific protein-DNA interactions were preserved. Interestingly, some cells displayed two MalI foci following Hex treatment, suggesting that plasmids, previously held together within the same condensate, were now free to diffuse apart (Supplementary Fig. S6A). We repeated this experiment with cephalexin-treated cells to increase the number of plasmids per cell (Supplementary Fig. S6B). We readily observed that, following Hex treatment, as ParB condensates dissolved, plasmids declustered. After removal of hexanediol, ParB condensates reformed efficiently and fused back, but MalI-*malO* foci often remained separate, presumably due to membrane anchoring (immobile foci). These experiments further support that coalescence is mediated by weak and/or non-specific interactions.

**Figure 5:**
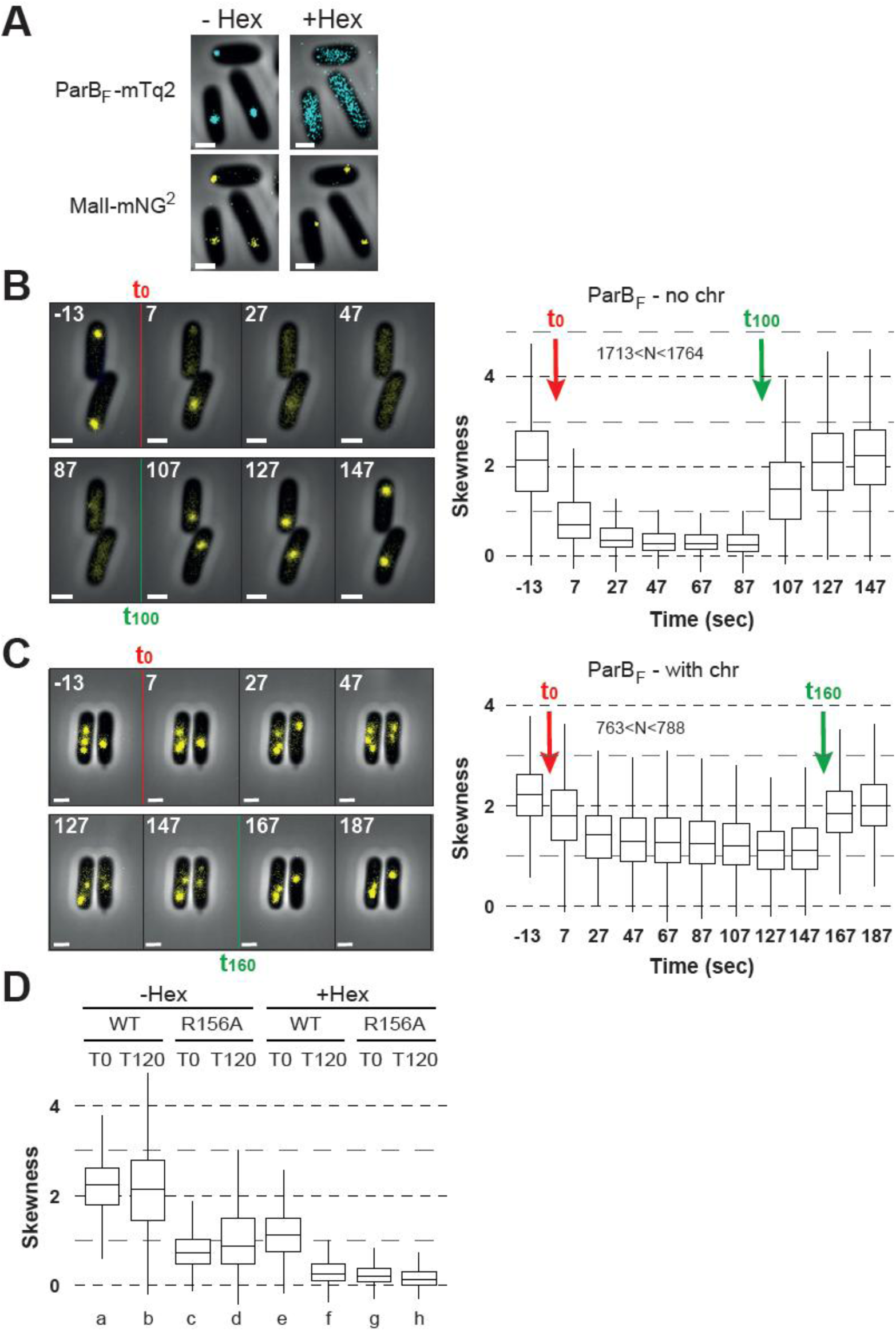
Assembly and disassembly of partition condensates occurs within seconds. **A**- Hexanediol disrupts ParB condensates but not DNA-bound proteins. Chromosome-depleted cells carrying the mini-F pPER08 were imaged before (-Hex; left panels) and after (+Hex; right panels) a 1-minute incubation with 10% 1,6-hexanediol in the blue (top panels) or yellow channels (bottom panels) to visualize ParB_F_-mTq2 or MalI-mNG^2^, respectively. **B**- Rapid and reversible disassembly of ParB condensates in nucleoid-free cells. (Left) Time-lapse imaging of chromosome-depleted cells expressing ParB_F_-mVenus from plasmid F was performed using a microfluidic setup, allowing controlled infusion of hexanediol at time 0 (T0) and wash out at 100 sec (T100). (Right) Quantification of ParB_F_-mVenus fluorescence skewness across > 1700 cells at each indicated time point. The rapid decrease in skewness upon Hex treatment and the fast recovery upon washout indicates a nearly instantaneous disassembly and reassembly of ParB condensates. **C**- Rapid, reversible but partial disassembly of ParB condensates in cells with an intact chromosome. Same as in (B) but performed in cells with a nucleoid. Hex was washed out after 160 s of infusion. Fluorescence skewness was calculated from > 760 cells per time point. **D**- Multiple interactions are involved in ParB condensate assembly. Fluorescence skewness was measured in cells expressing either ParB_F_ (WT) or the interaction-defective variant ParB_F_-R156A, from plasmid F, imaged before (T0) or after (T120) chromosome degradation, and either untreated (-Hex) or treated (+Hex) with hexanediol. Time-lapse microscopy was performed as in (B–C). The skewness levels in f, g and h are 0.33, 0.27 and 0.21, respectively.

To investigate the kinetics of ParB_F_ condensate disassembly, we performed time-lapse imaging during Hex infusion (Fig. 5B, top left). ParB_F_ fluorescence became visibly diffuse as early as 7 seconds after treatment and foci disappeared almost completely by 47 seconds. To quantify the Hex-induced disassembly, we measured the fluorescence skewness across 1150 cells (Fig. 5B, right). Before Hex addition, skewness was ∼2.11, slightly higher than measured in the presence of nucleoid (1.85; Fig. 2B). Upon Hex treatment, the average skewness progressively decreased to ∼0.84 and 0.47 after 7 and 27 seconds, respectively, reaching a plateau at ∼0.33 by 47 seconds. This plateau value is slightly above the skewness observed when all ParB_F_ are diffusive (0.19; Fig. 2B), consistent with only a small fraction of ParB_F_ remaining bound to *parS*_F_ or clamped nearby.

We then tested whether ParB_F_ condensates could reassemble after washing out Hex. Following a 100-seconds Hex exposure, cells were infused in buffer without Hex (Fig. 5B, bottom left). Within 7 seconds, skewness rapidly increased to 1.49, and by 27 seconds, it reached a value above 2, restoring the pre-treatment level and showing complete ParB_F_ condensates reassembly (Fig. 5B, right). Strikingly, the kinetics of reassembly mirrored that of disassembly, both occurring within seconds.

Altogether, these data demonstrate that ParB_F_ condensates disassemble and reassemble rapidly and reversibly in response to disruption and restoration of weak hydrophobic interactions, respectively. In the absence of the nucleoid, this behavior strongly suggests that the internal cohesion of partition condensates is primarily driven by reversible ParB_F_-ParB_F_ interactions.

### Partition condensates assembly requires both ParB_F_-ParB_F_ and ParB_F_-DNA interactions

To assess the relative contributions of ParB_F_-ParB_F_ and ParB_F_-nsDNA interactions to condensate assembly, we repeated hexanediol treatment experiments in cells with intact nucleoids (Fig. 5C). Under these conditions, condensate disassembly occurred slightly more slowly than in nucleoid-depleted cells, with skewness reaching a plateau within 47 seconds. Following Hex washout, ParB_F_ condensates reassembled almost completely within 7 seconds, indicating that the disruption and restoration of ParB_F_-ParB_F_ interactions proceed with slightly slower and faster kinetic, respectively, compared to nucleoid-depleted cells. Notably, while disassembly was nearly complete in nucleoid-depleted cells (plateau skewness ∼0.33), it was only partial in the presence of the chromosome, with skewness stabilizing at ∼1.2. These results indicate that, in addition to ParB_F_-ParB_F_ interactions, ParB_F_-DNA interactions substantially contribute to condensate integrity and assembly kinetics.

To further probe these interactions, we performed the hexanediol treatment in cells carrying a plasmid F encoding the ParB_F_-R156A variant (F1-10B15), which exhibits impaired ParB_F_-ParB_F_ interactions and reduced condensate formation (Fig. 2A, C). In nucleoid-depleted cells, ParB_F_-R156A-mVenus skewness dropped rapidly from 1.04 to 0.29 within 7 seconds, reaching a plateau at ∼0.21 (Supplementary Fig. S6C). This value closely matched that of fully diffuse ParB_F_ in the absence of condensates (∼0.19; Fig. 2B), indicating near-complete dissolution. In the presence of the nucleoid, skewness decreased from 0.81 to 0.36, plateauing at ∼0.28 (Supplementary Fig. S6D), slightly higher than in the nucleoid-free conditions but still indicative of near-total condensate disassembly.

Notably, in both WT and ParB_F_-R156A backgrounds, condensate disassembly was slower and less complete in the presence of the nucleoid than in its absence (Fig. 5B, C and Supplementary Fig. S6C, D). Conversely, condensate reassembly after hexanediol washout occurred slightly faster when the nucleoid was present. These kinetic differences underscore the contribution of ParB_F_-nucleoid interactions in both buffering condensates against chemically induced disruption and facilitating their rapid reformation.

A side-by-side comparison of WT and ParB_F_-R156A condensates across all conditions using skewness quantification (Fig. 5D) further supports that (i) condensates form in both nucleoid-containing and nucleoid-depleted cells (a-d), (ii) both hexanediol treatment and the ParB_F_-R156A variant cause partial defects in condensate formation (compare c and e to a), indicating that ParB_F_-ParB_F_ interactions are necessary but not sufficient for robust condensate assembly, and (iii) simultaneous nucleoid removal and perturbation of ParB_F_-ParB_F_ interactions (f) results in complete condensate disassembly.

Together, these findings establish that ParB_F_ condensate assembly is a cooperative process driven by both ParB_F_-ParB_F_ and ParB_F_-DNA interactions, with maximal cohesion and stability arising from their synergistic interplay.

## Discussion

### ParB condensates as a paradox of DNA segregation

In this study, we investigated the dynamics of ParB_F_ condensates involved in bacterial DNA partition, uncovering key kinetic and mechanistic features that define their behavior *in vivo*. Faithful segregation of low-copy-number genetic elements requires at least two partition complexes to remain physically separated on either side of the septum before cell division. This requirements presents a fundamental paradox: biomolecular condensates formed by phase separation inherently tend to coalesce into a single droplet to minimize surface tension and free energy (4). ParABS-mediated partition systems offer a powerful model for understanding how cells reconcile this conflict. Growing evidence indicates that ParB assemblies exhibit droplet-like properties both *in vivo* (9) and *in vitro* (11), making them well-suited to address how phase-separated condensates can remain stable as multiple entities inside a single bacterial cell.

### Chromosome removal reveals coalescence dynamics

Previous work showed that ParB_F_ condensates eventually merge *in vivo* following ParA_F_ depletion using a degron-based approach (9). However, fusion events became detectable only after overnight growth, due to continuous ParA_F_ overproduction followed by its gradual degradation, thereby precluding accurate analysis of coalescence kinetics. In the absence of ParA_F_, ParB_F_ condensates largely remain separated, likely because (i) their physical exclusion from the nucleoid confines them to the cell poles (22), limiting their probability of encounter and, (ii) they exhibit a reduced propensity to coalesce, consistent with a role for ParA_F_ in promoting ParB_F_ condensates assembly (see below).

To overcome these limitations, we developed a chromosome degradation approach that eliminates ParA_F_-dependent anchoring of ParB condensates to the nucleoid, thereby enabling their unrestricted diffusion in the cytoplasm. Under these conditions, nearly all ParB_F_ condensates coalesce within 120 minutes of chromosome removal (Fig. 1B-D and Supplementary Fig. S1), independently of their initial number (Fig. 3B). The resulting condensates exhibit high mobility, with diffusion coefficients reaching ∼0.02 and ∼0.04 µm² s^-1^ for F and mini-F plasmids, respectively, which is more than 10-fold higher than in nucleoid-containing cells (∼0.002 and ∼0.001 µm² s^-1^) (Fig. 4D and Supplementary Fig. S5). These values are comparable to diffusion rates reported for large nucleoprotein assemblies in *E. coli*, such as polysomes (∼0.04 µm² s^-1^), but remain lower than those of free ribosomal subunits (∼0.2 µm² s^-1^) (58,59). In nucleoid-free cells, the scaling exponent α derived from MSD analysis approaches ∼ 1, indicative of near-free Brownian motion, whereas in the presence of the chromosome, sub-diffusive values (α < 0.9) reflect constrained mobility. Together, these results strongly suggest that, in the absence of the nucleoid, the mobility of ParB_F_ condensates is primarily governed by their physical size, rather than by the length of the *parS*_F_-carrying plasmid.

Using transient splitting of ParB_F_ condensates as a reference point, we measured fusion times of ∼1 second (Fig. 4A-C). Such near-instantaneous coalescence upon contact indicates a droplet fusion process rather than Ostwald ripening (45). Moreover, the power-law distribution of coalescence times is consistent with first-encounter kinetics between diffusing particles, underscoring the stochastic nature of condensate collisions. Importantly, the increased mobility observed upon nucleoid removal directly enhances the encounter rate, thereby promoting rapid coalescence. These findings indicate that nucleoid tethering, mediated by ParA_F_, not only spatially organizes condensates but also kinetically limits their fusion by constraining their diffusion.

Strikingly, our analysis further suggests that ParB_F_ condensates may operate near the fusion-separation boundary (Supplementary Fig. S4), implying that only minimal energy is required either to separate fused droplets or to split condensates generated by partition complex duplication during replication. Operating at this boundary is likely crucial for DNA segregation, as it prevents irreversible coalescence into a single droplet while maintaining sufficient plasticity for dynamic reorganization. This finely tuned fusion-fission balance highlights how condensate phase behavior is optimized *in vivo* and raises the possibility that ParA_F_ may facilitate condensate separation with only a minimal energetic input. Alternatively, the initial separation step may occur independently of ParA_F_, as proposed for ParA from *C. crescentus* (60,61), where SMC (Structural Maintenance of Chromosome) proteins contribute to the separation of the replicated origins (62). In light of this finding, the role of ParAs in the separation and the positioning of duplicated partition, as well as other condensate-like cargos such as carboxysomes and chemotaxis clusters (13,63,64) warrants further investigation. Different cargos may have evolved distinct mechanisms to interact with their cognate ParA partners and to accommodate the specific biological constraints of each system, while still operating within the broader framework of the diffusion-ratchet model.

### ParA regulates both condensate positioning and assembly

Our results reveal that proper condensate assembly is essential for efficient coalescence. Perturbing the ability of ParB_F_ to form condensates (Fig. 2B), while preserving its capacity to bind to *parS*_F_ and diffuse nearby (36), strongly impaired coalescence, as observed with the ParB_F_-R156A variant (Fig. 2A). A similar reduction in coalescence was also observed in the absence of ParA_F_ (Fig. 2C). These observations suggest that ParA_F_ may contribute to the formation of a condensate-competent state of ParB_F_ *in vivo*, likely by stimulating ParB_F_-ParB_F_ interactions as reported *in vitro* (42). Supporting this hypothesis, ChIP-seq analyses have shown that ParB DNA-binding patterns around *parS* sites are strongly reduced in the absence of ParA, indicative of impaired higher-order assembly (65).

Our data reveal a dual role for ParA_F_ in regulating ParB condensate dynamics. First, ParA_F_ promotes the efficient assembly of ParB condensates, facilitating the cohesive interactions required for condensate fusion. Second, through its well-established function in tethering partition complexes to the nucleoid (22), ParA_F_ spatially separates condensates, thereby limiting their encounters and preventing uncontrolled fusion. This dual functionality allows ParA_F_ to simultaneously enhance condensate fusion competence while reducing the likelihood of fusion events. In the absence of ParA_F_, spatial constraints that normally maintain condensate separation are relaxed, yet the capacity for fusion is also reduced due to impaired condensate assembly. Thus, ParA_F_ acts as a balanced coordinator, ensuring that partition complexes are both properly assembled and correctly positioned within the nucleoid, fine-tuning their dynamics to ensure faithful DNA segregation.

### Rapid kinetics of ParB condensates assembly

ParB condensates associated with plasmids or chromosomes are continuously present in cells, provided that at least one cognate *parS* site is present (12). Disassembly and reassembly of the ParB_F_ complex have been reported to occur within less than a minute during replication of the *parS*_F_ centromere region (66). In chromosome-depleted cells treated with 1,6-hexanediol, which disrupts weak hydrophobic interactions (57), we find that these kinetics are even faster, occurring within only a few seconds, with most ParB_F_ dispersing over the nucleoid (Fig. 5A-B). This treatment, intended to disrupt interactions between ParB dimers, may also interfere with the *parS*- and CTP-dependent intra-dimer association of the two N-terminal domains of ParB_F_ (30,32), thereby promoting clamp opening and condensates disassembly. It may also perturb ParA_F_-ParB_F_ interactions which are shown to promote ParB_F_-ParB_F_ interactions *in vitro* (42), further contributing to condensate disassembly.

Importantly, this disassembly is fully and rapidly reversible, as ParB_F_ condensates reassemble within seconds following hexanediol removal, restoring initial fluorescence skewness levels (Fig. 5B). Such rapid re-assembly is consistent with the high dynamics observed in ParB complexes (24,31,41) and with the exchangeable nature of biomolecular condensates (5). Alternatively, models based on sequential ParB loading at *parS*, followed by CTP-dependent clamp formation and sliding to mediate long-range DNA bridging (31,38,41), would require very high turnover rates. Such rate would be necessary to account for the rapid conversion of large numbers of ParB molecules into a condensed state and to sustain a nodal accumulation.

To date, only a single exception to the continuous presence of ParB condensates has been reported. In the predatory bacterium *Bdellovibrio bacteriovorus*, ParB*_Bbac_* condensates are absent during a defined developmental stage despite expression of ParB*_Bba_* (67). The molecular basis of this stage-specific suppression of condensate formation remains unknown, but it correlates with the presence of short DNA segments adjacent to the two *parS* sites. This observation suggests that the initial binding step at *parS_Bbac_* may be inhibited by specific DNA context or structure, and possibly regulated through the action of a developmentally controlled factor to prevent ParB condensate assembly. Together, these observations support the view that while ParB condensates can assemble extremely rapidly once nucleation occurs at *parS*, modulation of this nucleation step may provide an efficient mechanism to regulate condensate formation in specific cellular contexts.

### Mechanistic model of nucleation and stabilization

In nucleoid-containing cells, hexanediol treatment induced only partial disassembly of ParB_F_ condensates, with fluorescence skewness plateauing at ∼1.2 versus ∼0.33 in nucleoid-free cells (Fig. 5B-C). This difference indicates that interactions with chromosomal DNA, most likely through non-specific binding, contribute to stabilizing ParB condensates and buffer them against complete dispersion. Such behavior is more consistent with a phase separation mechanism than with a model based solely on long-range ParB bridging interactions. In a purely bridging-based model (10,41), disruption of ParB-ParB interactions between clamps or the release of ParB clamps from *parS*-proximal DNA would be expected to yield comparable dispersion of ParB regardless of the presence of the chromosome. The pronounced difference observed between nucleoid-containing and nucleoid-depleted cells therefore argues for an additional stabilizing contribution arising from ParB-nsDNA interactions within the nucleoid environment. An alternative, non-exclusive possibility is that a subset of ParB clamps is loaded *in trans* onto chromosomal DNA, as reported *in vitro* for ParB*_Bsub_* (38). In such a scenario, chromosome removal would release these ParB, allowing unrestricted diffusion, whereas in nucleoid-containing cells they would remain spatially constrained. However, this model would require a substantial fraction of ParB_F_ to be loaded *in trans* relative to *parS*_F_, a possibility that remains to be directly tested *in vivo*.

Consistent with a central role for ParB multimerization, the ParB_F_-R156A variant, defective in ParB_F_-ParB_F_ interactions but still competent for clamp formation at *parS*_F_ (36), undergoes near-complete dissolution both in the presence or absence of chromosome (Fig. 5D, Supplementary Fig. S6C-D). This result underscores the primary role of ParB_F_ multimerization in condensate formation, while ParB_F_-DNA interactions provide an additional stabilizing layer. Interestingly, the residual skewness observed in the presence of hexanediol (0.33) remains higher than that measured in cell lacking *parS*_F_ (∼0.19; Fig. 2B), suggesting that a fraction of ParB_F_ dimers remains associated with *parS*_F_ or persists as sliding clamps on *parS*_F_-flanking DNA. These residual, potentially activated ParBs may serve as nucleation seeds, enabling rapid and efficient reassembly of condensates once hydrophobic interactions are restored.

Altogether, these observations support a model in which ParB condensates emerge from cooperative ParB-ParB and ParB-DNA interactions. ParB bound to *parS* and subsequently converted into clamps on flanking DNA likely serves as a nucleation site for droplet formation. These clamped ParB species would function as activation seeds that recruit diffusive ParB, enabling the rapid and reversible condensate dynamics observed here. This mechanistic model aligns well with our recent proposal suggesting that clamped ParB species are activated intermediates promoting condensate formation (36).

## Conclusion

In summary, our findings further support the view that ParB_F_ partition complexes are *bona fide* biomolecular condensates. They undergo rapid, reversible fusion and fission, display high mobility when untethered, and rely on multivalent interactions for assembly. We further show that ParA_F_ acts not only as a spatial positioning factor but also as a key regulator of condensate coalescence. By preventing uncontrolled fusion of partition condensates while promoting their even spatial distribution, ParA_F_ ensures that multiple partition complexes remain as distinct entities within the same cell, thereby securing faithful DNA segregation.

More broadly, our results provide a biophysical framework for understanding bacterial DNA segregation as a process governed by regulated phase separation and ParA-type ATPase dynamics. Partition condensates exemplify how bacterial cells can exploit the physical principles of phase separation to achieve robust and controlled spatial organization. This work reinforces the growing concept that phase separation represents a conserved and versatile strategy for organizing membrane-less compartments across both prokaryotic and eukaryotic systems.

## Acknowledgements

We are grateful to Julien Cayron and Christian Lesterlin for sharing the strain LY463 prior to publication, and to Johann Paulson for the gift of plasmid pRLT1. We are grateful to Jian Liu and Manuel Campos for critical reading of the manuscript, and we thank Christian Lesterlin, Anthony Vecchiarelli and Fabian Erdel for insightful discussions and suggestions. We also thank all members of the GeDy (CBI) and the SCPN (L2C) teams for fruitful discussions.

## Authors contributions

Conceptualization, J.-Y.B.; Data curation, P.R., J.-Y.B.; Formal analysis, L.D., J.C., J.-Y.B., and J.-C.W.; Funding Acquisition, J.-Y.B.; Investigation, P.R., L.D., J.C., J.R., J.-C.W., and J.-Y.B.; Methodology, P.R., L.D., J.R., J.-C.W., and J.-Y.B.; Project administration, J.-Y.B.; Resources, F.C., J.-C.W., J.-Y.B.; Supervision, J.-C.W., J.-Y.B.; Validation, P.R., L.D., J.-C.W., and J.-Y.B.; Visualization, P.R., J.-C.W. and J.-Y.B.; Writing – Original draft, P.R., L.D., F.C., J.-C.W. and J.-Y.B.; Writing – Review & Editing, P.R., L.D., F.C., J.-C.W., and J.-Y.B.

## Data availability

All data generated or analyzed during this study are included in this published article and its supplementary information files.

## Ethics statement

This article does not contain any studies with human or animal subjects.

