## Supplementary material for "Fast assembly and *in vivo* coalescence of ParB_F_ biocondensates involved in bacterial DNA partition": Supplemenraty data

**The PDF file includes:**

- Supplementary Figures S1 to S6
- Supplementary Tables S1 to S2
- Supplementary Materials and methods
- Supplementary References

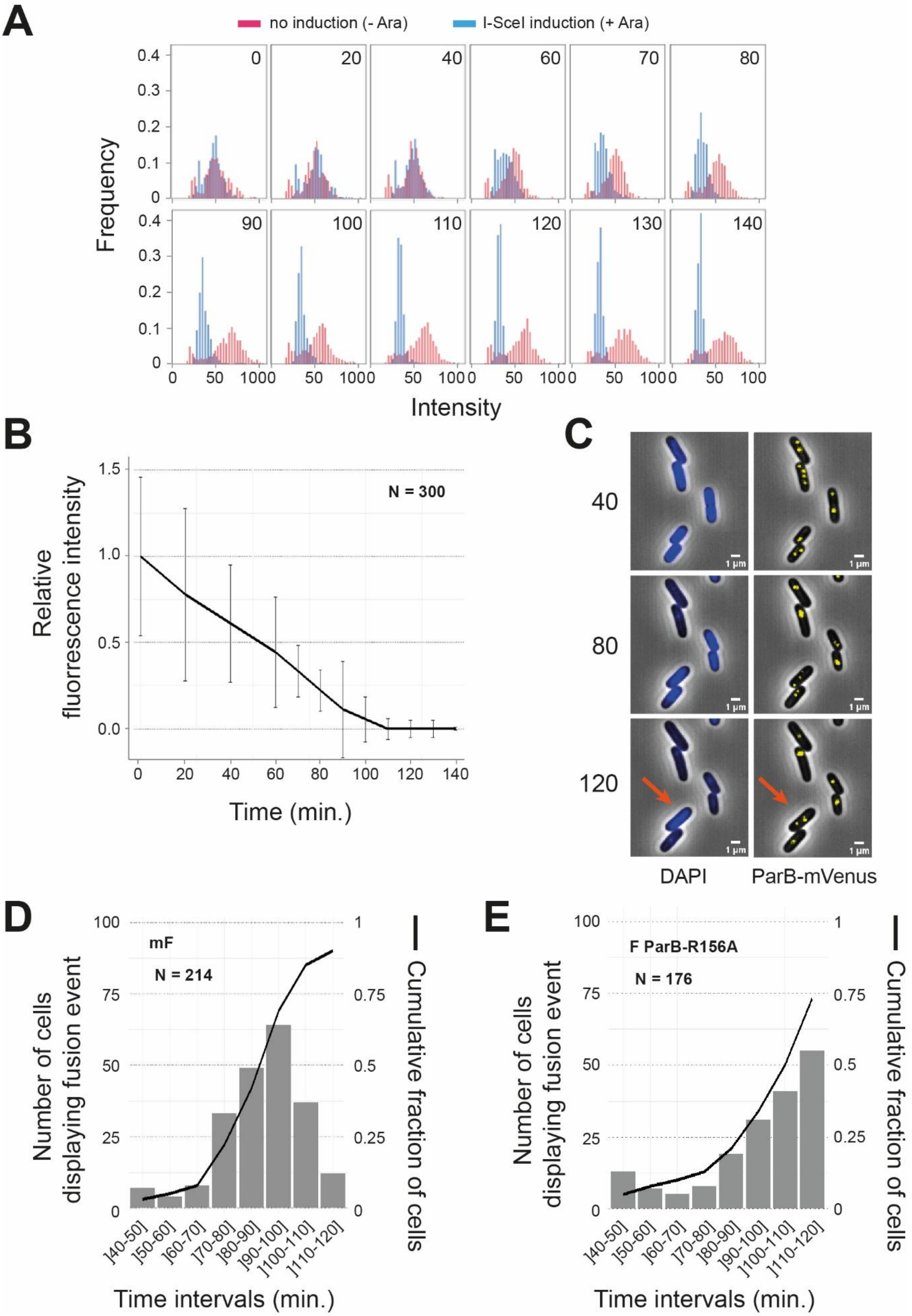

A
no induction (- Ara)
I-SceI induction (+ Ara)
Frequency
Intensity
0 20 40 60 70 80 90 100 110 120 130 140
B
Relative fluorescence intensity
N = 300
Time (min.)
C
40
80
120
DAPI
ParB-mVenus
1 µm
D
mF
N = 214
Number of cells displaying fusion event
Cumulative fraction of cells
Time intervals (min.)
]40-50] ]50-60] ]60-70] ]70-80] ]80-90] ]90-100] ]100-110] ]110-120]
E
F ParB-R156A
N = 176
Number of cells displaying fusion event
Cumulative fraction of cells
Time intervals (min.)

**Supplementary Figure S1:** Chromosome degradation induces ParB condensate fusion.
**A-** Time course of chromosome degradation monitored by DAPI staining under fluorescence microcopy. Strain DLT4156 carrying pSN1 was grown to exponential phase and either induced with arabinose (blue bars) to express I-SceI or left uninduced (red bars). At indicated time points (0-140 min), cells were stained with DAPI, and fluorescence intensity per cell was quantified. Distributions of DAPI intensity are shown for $N$ = 300 cells per condition and time point.
**B-** Quantification of chromosome degradation over time. Average DAPI fluorescence intensity per cell from panel A was plotted over time, showing progressive DNA loss following arabinose induction. $N$ = 300 cells.
**C-** Representative images showing simultaneous chromosome degradation and ParB condensate dynamics. Cells were stained with DAPI and imaged at 40, 80, and 120 min post-induction in both blue (DAPI; left panels) and yellow (ParB-mVenus; right panels) channels. The red arrow highlights a cell in which the chromosome remains partially intact and in which ParB condensates have not fused. Scale bar: 1 µm.
**D-** Quantification of ParB-mVenus fusion events over time for mini-F (pJYB234). Time-lapse imaging was performed every 2 minutes between 40 and 120 min post-induction. For each time point, the number of cells displaying a fusion event was recorded. The solid line indicates the cumulative fraction of cells showing both a fusion event and complete chromosome degradation. $N$ = 214 cells.
**E-** Same as in D, but for the full-length F plasmid expressing the $ParB_F$-R156A variant (F1-10B08). $N$ = 176 cells.

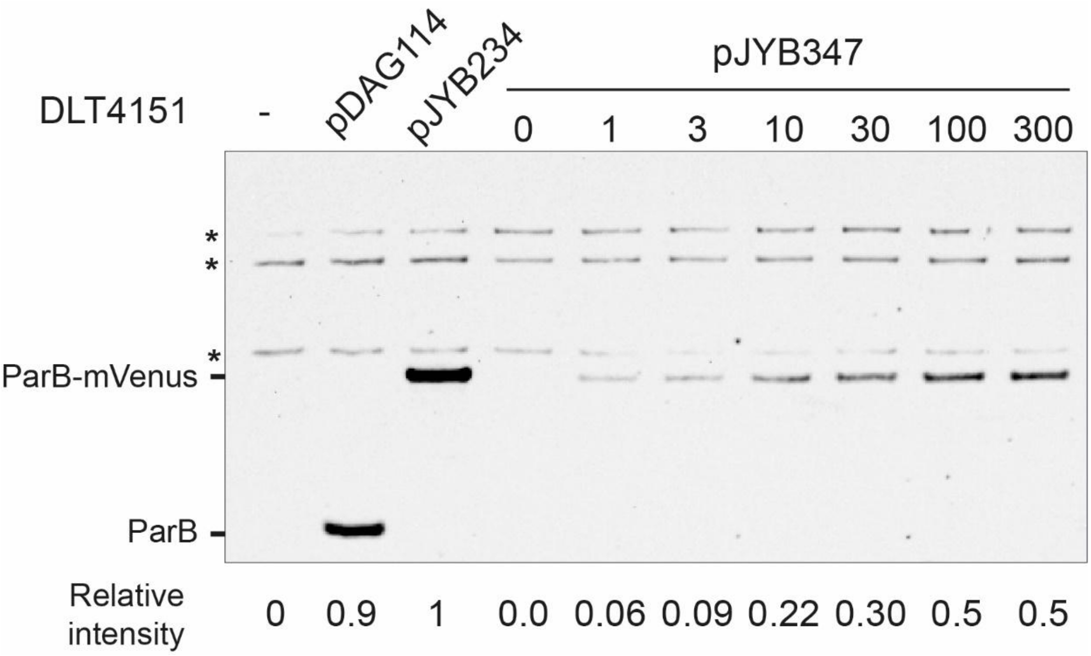

**Supplementary Figure S2:** Expression level of $ParB_F$-mVenus.
A typical Western blot using anti ParB antibodies is displayed for the strain DLT4151, carrying or not (-) the indicated plasmids, grown to exponential phase. For strain carrying pJYB347, anhydrotetracycline was added at the indicated concentrations (in µg.ml$^{-1}$) to induce expression from the *tet* promoter. Relative levels of $ParB_F$ or $ParB_F$-mVenus (compared to expression from pJYB234, set as 1) are shown below each lane. Signal intensities were normalized using internal loading controls, based on cross-reacting bands marked with asterisks (*).

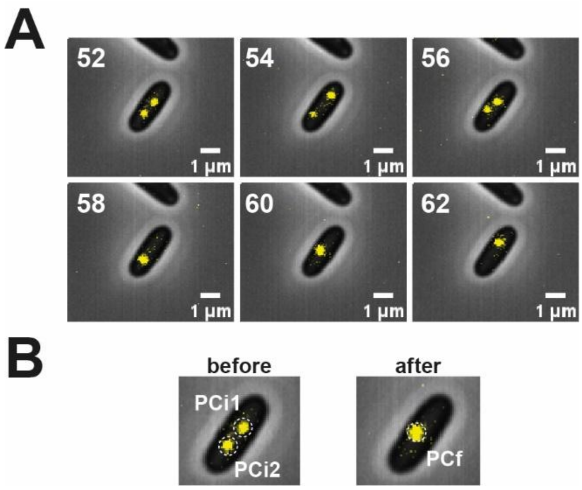

**Supplementary Figure S3:** Imaging and quantification of ParB condensate coalescence.
**A-** Time lapse imaging of $ParB_F$-mVenus condensate fusion. Images were acquired every two minutes between 40 to 120 minutes post-arabinose induction. Representative successive images are shown before and after a fusion event at the indicated time point (min). Scale bar = 1 µm.
**B-** Quantification of ParB condensate fluorescence intensity. Regions of interest (ROI) were manually drawn around ParB foci to measure fluorescence before and after fusion. The summed intensity of the two individual foci (PCi1 + PCi2) was average over three successive images prior to fusion and compared to the average fluorescence intensity of the resulting single focus (PCf), also average over three successive images. This ratio was used to calculate the normalized fluorescence change displayed in Fig. 3A.

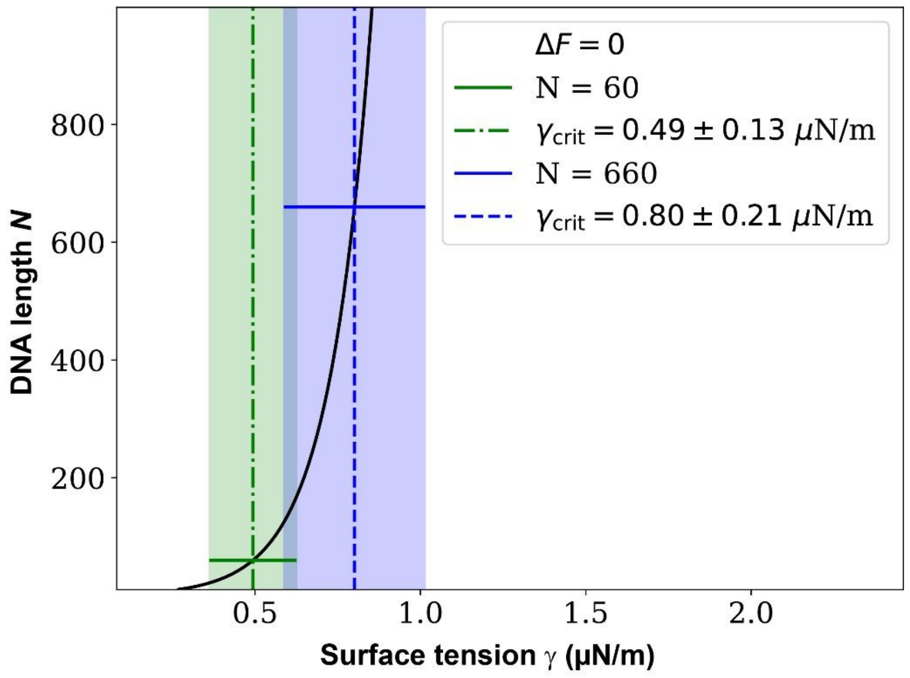

**Supplementary Figure S4:** Phase diagram of the fusion and splitting transition of ParB-DNA. The phase diagram is plotted in the plane (γ, *N*), where γ is the surface tension of the condensates and *N* the length of the plasmid in monomers. Here one monomer corresponds to 50 nm (*i.e.*, ~150 bp). The black curve represents the boundary where the free-energy difference between fused and split condensates is null ($\Delta F = 0$), separating regimes where fusion (right) or splitting (left) is thermodynamically favored (see calculation in Supplementary Methods). Solid horizontal lines indicate polymer lengths corresponding to the mini-F (green, $N = 60$) and full-length F (blue, $N = 660$) plasmids. Dashed vertical lines mark the critical surface tensions $\gamma_{crit} \pm \sigma$ determined for each plasmid (mini-F: $0.49 \pm 0.13$ μN.m$^{-1}$, green; F: $0.80 \pm 0.21$ μN.m$^{-1}$, blue), with shaded bands indicating their uncertainties.

Note: In the absence of the chromosome, the $ParB_F$-R156A variant effectively reduces the $ParB_F$-$ParB_F$ interaction strength within this framework. This reduction corresponds to a decrease in the effective surface tension γ, shifting the system toward the low-γ regime. Under these conditions, coalescence is no longer thermodynamically favored, resulting in the formation of multiple, weakly interacting condensates. These condensates are expected to be only marginally stable, or even transient, placing the system near the boundary of the framework's applicability.

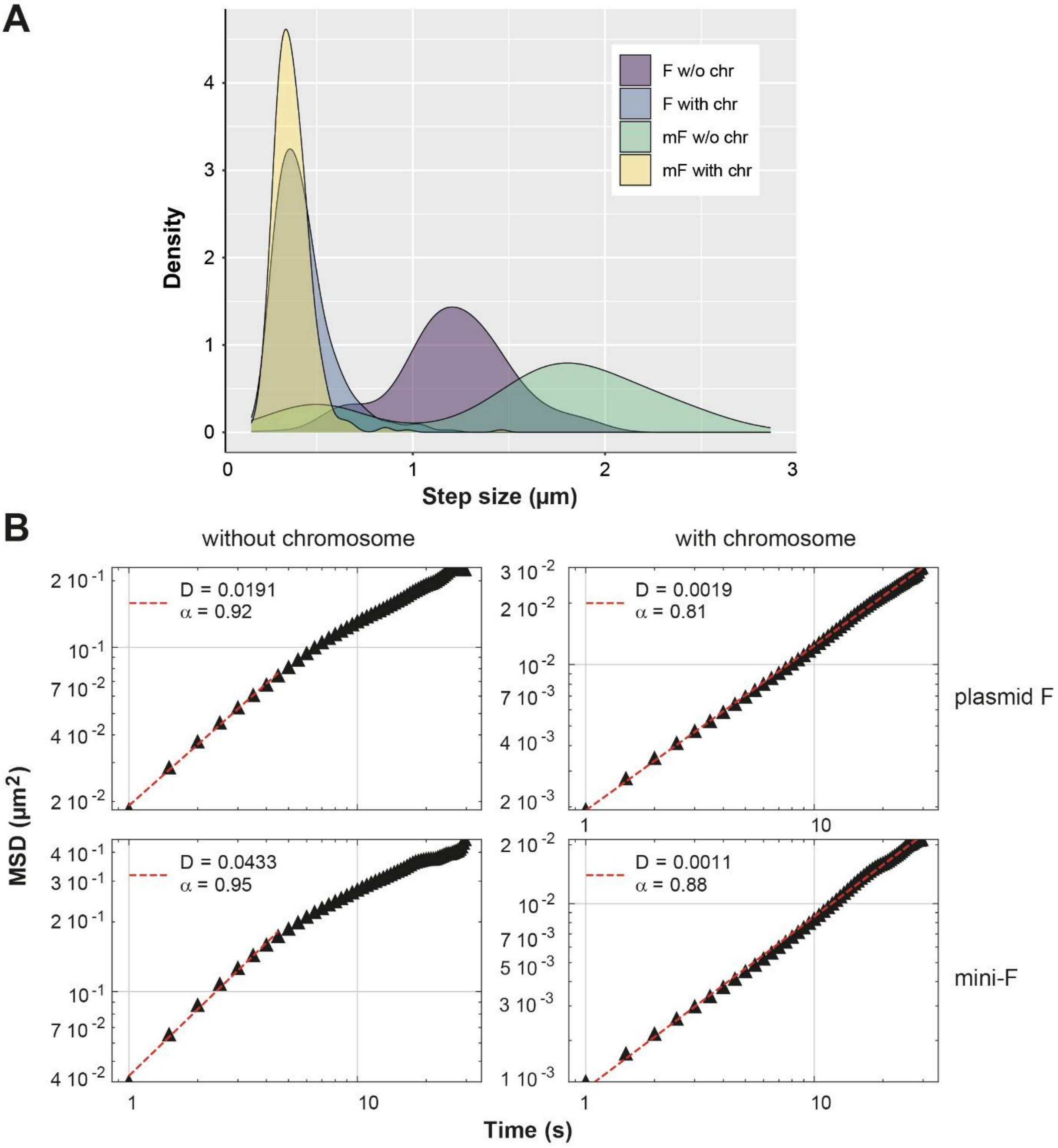

**Supplementary Figure S5:** Mobility of ParB condensates in the presence and absence of chromosome.

ParB$_F$-mVenus foci from both plasmid F and mini-F were tracked at 0.5 second intervals over a 30 seconds period in cells with or without a chromosome. Quantification was performed on 707 tracks (with chromosome) and 230 tracks (without chromosome) for plasmid F, and on 427 tracks (with chromosome) and 274 tracks (without chromosome) for mini-F.

**A-** Step size distributions reveal increased mobility in the absence of chromosome. The distribution of step sizes between successive frames is shown in violet and blue for plasmid F in the presence (with) and absence (w/o) of the chromosome, respectively, and in green and yellow for mini-F in the presence (with) and absence (w/o) of the chromosome, respectively.

**B-** Mean squared displacement (MSD) and diffusion coefficients. MSD analyses were performed for ParB$_F$ foci from plasmid F (top panels) and mini-F (bottom panels), in the absence (left panels) or presence (right panels) of the chromosome. For each condition, the diffusion coefficient (D, in $\mu m^2.s^{-\alpha}$) and the MSD scaling exponent ($\alpha$) are indicated.

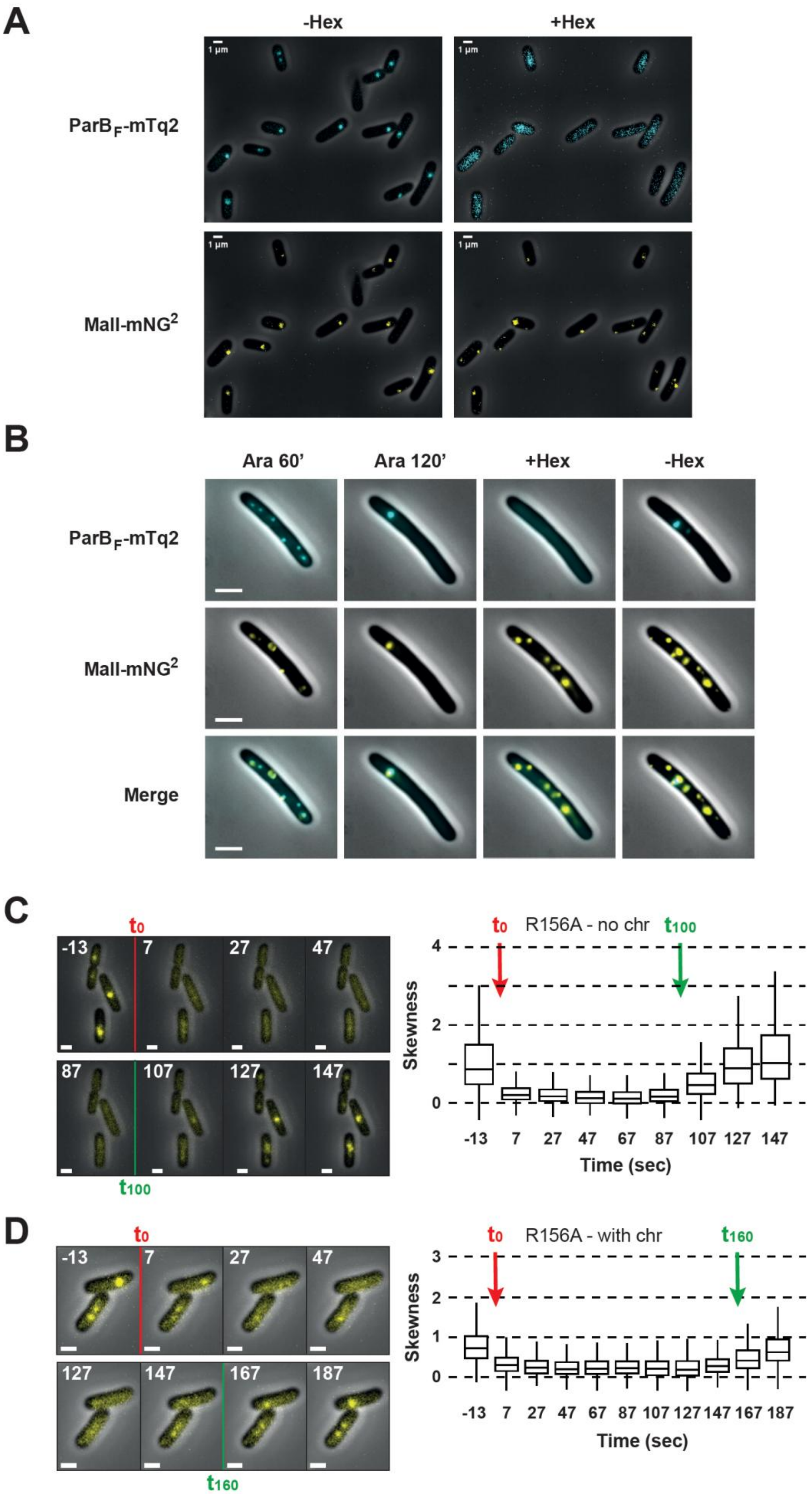
A
-Hex
+Hex
ParB$_F$-mTq2
MalI-mNG$^2$
1 µm
B
Ara 60'
Ara 120'
+Hex
-Hex
ParB$_F$-mTq2
MalI-mNG$^2$
Merge
C
$t_0$
$t_{100}$
-13
7
27
47
87
107
127
147
R156A - no chr
Skewness
Time (sec)
D
$t_0$
$t_{160}$
-13
7
27
47
127
147
167
187
R156A - with chr
Skewness
Time (sec)

**Supplementary Figure S6:** Hexanediol specifically disassembles ParB condensates.

**A-** Hexanediol disrupts ParB condensates but not DNA-bound proteins. Chromosome-depleted cells (DLT4234) carrying the mini-F plasmid pPER08 were imaged before (-Hex; left panels) and after (+Hex; right panels) a 1-minute incubation with 10% 1,6-hexanediol. Images show $ParB_F$-mTq2 fluorescence (blue channel; top panels) and MalI-$mNG^2$ bound to a 7×*malO* array (yellow channel; bottom panels). ParB condensates rapidly dispersed upon Hex treatment, whereas MalI foci remained unaffected, showing that hexanediol selectively disrupts ParB condensates while preserving stable DNA-protein interactions.

**B-** Plasmids decluster concomitantly to ParB condensate dissolution upon hexanediol treatment. DLT4234 cells were grown to exponential phase, treated with cephalexin 30 min., and further incubated with cephalexin and arabinose (Ara) for 120 min. Hexanediol was then added for 5 min. before wash-out. Images were capture at 60 min. (Ara 60') and 120 min. (Ara 120') after addition of Ara, and 2 min. after addition of Hexanediol (+Hex) and 2 min. after wah-out (-Hex). Fluorescence was visualized in the blue channel ($ParB_F$-mTq2; top), yellow channel (MalI-$mNG^2$; middle), and merged (bottom). Note that, after the wash-out step (-Hex), the $mNG^2$ foci become mostly immobile, likely due to membrane tethering.

**C-** Rapid and reversible disassembly of $ParB_F$-R156A-mVenus condensates in nucleoid-free cells. (left) Time-lapse imaging of chromosome-depleted cells expressing $ParB_F$-R156A-mVenus from plasmid F was performed using a microfluidic setup, allowing controlled infusion of hexanediol at time 0 (T0) and wash out at 100 sec (T100). (right) Fluorescence skewness was quantified from > 1082 cells at each time point and highlights the sharp drop in skewness followed by rapid recovery upon washout.

**D-** Same as (B), but in cells with an intact chromosome. Hexanediol was washed out after 160 s of infusion. Fluorescence skewness was calculated from > 845 cells per time point.

**Supplementary Table S1:** Bacterial strains and plasmids.

| *Strain* | *Genotype/Relevant properties* | *Source/Reference* |
|---|---|---|
| DLT1215 | W1485 *thi leu thyA deoB supE* Δ*(ara-leu)7696 zac3051::Tn10 rpsL812* | (Bouet et al., 2007) |
| DLT3258 | DY378 / pAM238 | This work |
| DLT3589 | DLT1215 / F1-10B04 | (Debaugny et al., 2018) |
| DLT3712 | DLT3258 / F1-10B | This work |
| DLT3713 | DLT3258 / F1-10B04 | This work |
| DLT4108 | DLT1215 / pCP20 | This work |
| DLT4114 | DLT1215 / F1-10B15 | This work |
| DLT4149 | LY643 *kan-Pcp18::araE533* | This work |
| DLT4151 | DLT4149 *srlC::tn10, recA* | This work |
| DLT4156 | DLT4151 / F1-10B04 | This work |
| DLT4161 | DLT4151 / pJYB234 | This work |
| DLT4199 | DLT4151 / pJYB347 | This work |
| DLT4214 | DLT4151 / F1-10B15 | This work |
| DLT4215 | DLT4151 / F1-10B12 | This work |
| DLT4222 | DLT4151 / pPER07 | This work |
| DLT4234 | DLT4151 / pPER08 | This work |
| DLT4378 | DLT4151 / pPER16 | This work |
| DY378 | *lacI857* Δ*(cro-bio)* | (Yu et al., 2000) |
| LY643 | MG1655 *TB28 ilvA::ISce1CS-frt, codA::ISce1CS-frt, yeiU::ISce1CS-frt, ydeO::ISce1CS-frt* | Gift from C. Lesterlin (to be described elsewhere) |

| *Plasmid* | *Relevant characteristics* | *Source/Reference* |
|---|---|---|
| F1-10B | F1-10 *ccdB cat*$^+$ | (Debaugny et al., 2018) |
| F1-10B04 | F1-10B *parB*$_F$*-mVenus* | (Debaugny et al., 2018) |
| F1-10B12 | F1-10B04 Δ*parA PltetO-ParB-mVenus* | This work |
| F1-10B15 | F1-10B *parB*$_F$*-R156A-mVenus* | This work |
| pAM238 | pGB2 derivative, spectimomycin resistant | (Bouet et al., 1996) |
| pCP20 | pSC101 *rep*$^{ts}$, *flp* | (Datsenko and Wanner, 2000) |
| pDAG114 | mini-F *repFIA*$^+$, *ccdB*$^-$, *resD*$^+$, *rsfF*$^+$, *cat*$^+$ , *parABS*$_F^+$ | (Lemonnier et al., 2000) |
| pDAG198 | pDAG114 Δ*parA PltetO-parB* | (Castaing et al., 2008) |
| pDAG209 | pDAG114 Δ*parAB, parS*$_F^+$ | (Bouet et al., 2006) |
| pJYB212 | pDAG114 *parB*$_F$*-mEos2* | (Sanchez et al., 2015) |
| pJYB234 | pDAG114 *parB*$_F$*-mVenus* | (Diaz et al., 2015) |
| pJYB240 | pJYB212 *parB*$_F$*-mTurquoise2* | This work |
| pJYB342 | pDAG114 *parB*$_F$*_R156A* | (Delimi et al., 2025) |
| pJYB347 | pDAG198 Δ*parA PltetO-ParB-mVenus* | This work |
| pKD4 | R6K *FRT-kan-FRT* | (Datsenko and Wanner, 2000) |
| pPER07 | pDAG209 *malO/malI-mNG*$^2$ * | This work |
| pPER08 | pJYB240 *malO/malI-mNG*$^2$ * | This work |
| pPER16 | pJYB234 *ΔparS*$_F$ | This work |
| pRLT1 | *malO/malI-mNG*$^2$ * | Gift from J. Paulsson |
| pSN1 | P$_{BAD}$::I-Sce1 | (Stracy et al., 2021) |

* *malI-mNG*$^2$ stands for the fusion of a tandem dimer of *mNeonGreen* to *malI: malI-mNeonGreen-mNeonGreen*

**Supplementary Table S2:** List of oligonucleotides. The sequences are displayed in 5′ to 3′ orientation.

| *Name* | *Sequence* |
|---|---|
| JRP01 | CTCGCCAGTTCGCTCGCTATGCTCGG |
| JRP02 | GCGCAAAATGTTGTGGATAAGCAGG |
| JRP03 | CCTGCTTATCCACAACATTTTGCGCCCATGGTCCATATGAATATCCTCC |
| JRP04 | GATTGTGTAGGCTGGAGCTGCTTC |
| JRP05 | GAAGCAGCTCCAGCCTACACAATCGTGCCGGCACGTTAACCGGGCTGC |
| JRP06 | GATACAACGCCACGCCCAGCAGCCG |
| JRP07 | CGTTTTCAGGCCGTCATCGAGGAAGCGTTCGGCTAGGGCGGCGTAGCACCAGGCG TTTAAGGGCACC |
| JRP08 | GGAGGTGTTCGGCATACATCTGATC |
| JRP09 | AACAGTACTGCGATGAGTGGCAGGG |
| JRP10 | GCCTGTCAAGGGCAAGTATTGACATGTC |
| JRP11 | AGGCGCACGCTTCATGGGCCCGGTACCTTTCTCCTCTTTA |
| JRP12 | ACGGATTCACCGTTTTTATCAGGCTCTG |
| JRP13 | ATGAAGCGTGCGCCTGTTATTCCAAAAC |
| PR06 | AAACGGTGAATCCGTTAGCGAGGTG |
| PR11 | GTACACGACGTCAATTGTGAGCGGATAACAATTTCAC |
| PR18 | TACACCACGCGTAATTGTTATCCGCTCACAATTCTAGACGCC |
| PR19 | GTACACGACGTCTAGCGGTGCGTCCCTGTTTGCATTATG |
| PR20 | TACACCACGCGTACGCAGGTTAAGCTGGCTTAGCATC |
| PR21 | CCCGCCATCACCAGGCCTGATGGG |
| PR22 | AGCTGAAGCTCGAATTTCGCCGCCC |
| PR25 | CCCATCAGGCCTGGTGATGGCGGGGCTCGAGGAATTCTCCAGATTCTAG |
| PR26 | GGGCGGCGAAATTCGAGCTTCAGCTGAGCTCTTACTTGTACAATTCGTCC |

### Supplementary Materials and methods

#### Strain constructions

All strains used in this study are derivatives of *E. coli* K12 and are listed, along with plasmids, in Table S1.

Strain LY643, a derivative of TB28 (Lesterlin et al., 2014) carrying four I-SceI sites, was constructed by Julien Cayron (gift from C. Lesterlin) and will be described elsewhere. Strain DLT4149 was obtained by transducing the *Frt-kan-Frt-Pcp18::araE* locus from DLT2200 (our laboratory collection) into LY643. This strain was subsequently modified by transduction of *recA1*, *srlC*::*tn10* from JS238 (team collection), yielding DLT4151.

#### Plasmid constructions

All plasmids used in this study are listed in Table S1. The oligonucleotides and their sequences are provided in Table S2.

pJYB347 was derived from pDAG198 by inverse PCR using primers JRP11 and JRP12 (see Table S2). A DNA fragment containing *parB*$_F$*-mVenus* and *parS*$_F$ was PCR-amplified from pJYB234 (Diaz *et al.*, 2015) using primers JRP13 and PR06, and inserted into the pDAG198 backbone by the Gibson assembly (Telesis Bio).

F1-10B12 was derived from F1-10B (Debaugny et al., 2018). A DNA fragment containing *Ptet_ΔparA*$_F$*_parB*$_F$*-mVenus_parS*$_F$ was PCR-amplified from pJYB347 using primers JRP02 and JRP07. Sequential PCRs were then performed with primers pairs JRP08/JRP09 and JRP06/JRP10 to add homology arms for recombination with F1-10B. The final PCR product was purified and transformed in DLT3712, a DY378 strain conjugated with F1-10B, and recombination was achieved via λ-RED recombineering (Datsenko and Wanner, 2000). The recombinant F1-10B was conjugated into DLT1215 carrying pCP20 to excise the *FRT-kan-FRTI* cassette. pCP20 was then cured by incubation at 42°C, yielding strain DLT4215 carrying F1-10B12.

F1-10B15 was derived from F1-10B04 (Debaugny et al., 2018). The insert was assembled via three overlapping PCR steps. First, *parB*$_F$*_R156A-mVenus* was amplified from pJYB342 using primers JRP01 and JRP02. Next, a kanamycin resistance cassette was amplified from pKD4 using primers JRP03 and JRP04. Lastly, homology to the plasmid F was added by amplification from F1-10B04 using primers JRP05 and JRP06. The final PCR product was purified and processed as indicated for F1-10B12, resulting in stain DLT1215 carrying F1-10B15.

pPER07 was derived from pDAG209 by inverse PCR using primers PR21 and PR22. A DNA fragment containing the FROS MalI-mNG$^2$/*malO* system was amplified from pRLT1 using primers PR25 and PR26, which included regions homologous to the ends of the PCR-amplified pDAG209 fragment. Both fragments were co-transformed into *E. coli* DH5α and the recombinant plasmid was obtained by homologous recombination.

pJYB240 was constructed from pJYB212 by replacing the *mEos2* gene with a PCR-amplified mTurquoise2 fragment (Goedhart et al., 2012) using the In-Fusion cloning kit (Clontech). The fluorescent protein was fused in-frame with *parB*$_F$.
Plasmid pPER08 was generated from pJYB240 by inverse PCR using primers PR19 and PR20, which carried floating tails with *Aat*II and *Mlu*I restriction sites, respectively. The FROS MalI-mNG$^2$/*malO* cassette was amplified from pRLT1 using primers PR11 and PR18, which also contained *Aat*II and *Mlu*I sites. Both the vector and insert were digested with *AatI*I and *Mlu*I, and ligated using T4 DNA ligase (New England Biolabs).

pPER16 was derived from pJYB234 by inverse PCR using phosphorylated primers PR19 and PR20, flanking the *parS*$_F$ site. The PCR product was purified and self-ligated, resulting in a precise deletion of the *parS*$_F$ site.

All mini-F plasmids were sequence-verified.

**Mean Squared Displacement (MSD) analyzes**

Each individual track consists of $N = 60$ measures of the position of the complex, spaced by a constant time step $\delta t = 0.5s$. The positions are $\vec{R}(t)$ where $t = 0, 1, \ldots, N-1$. The MSD is defined (for lag time $\Delta t = 0, 1, ..., N-1$) as:

$$\mathrm{MSD}(\Delta t) = \frac{1}{N-\Delta t} \sum_{t=0}^{N-\Delta t-1} [\vec{R}(t+\Delta t) - \vec{R}(t)]^2 \quad (1)$$

$$= \frac{1}{N-\Delta t} \left( \underbrace{\sum_{t=0}^{N-\Delta t-1} R^2(t+\Delta t) + R^2(t)}_{A(\Delta t)} -2 \underbrace{\sum_{t=0}^{N-\Delta t-1} \vec{R}(t+\Delta t) \cdot \vec{R}(t)}_{B(\Delta t)} \right) \quad (2)$$

To speed up MSD calculation, we use a fast correlation algorithm (Calandrini *et al.*, 2011).

1. $A(\Delta t)$ is calculated throught a recursive expression :

$$A(\Delta t + 1) = A(\Delta t) - R^2(\Delta t) - R^2(N_t - \Delta t - 1) \quad (3)$$

$$A(0) = 2 \sum_{t=0}^{N_t - 1} R^2(t) \quad (4)$$

2. $B(\Delta t)$ is similar to the autocorrelation function, and thanks to the Wiener-Khinchin theorem we can speed up its calcuation using the Fast Fourier Transform (FFT).

Furthermore, we performed a Principal Component Analysis (PCA) to determine a principal direction along which we project the positions. In conditions where the nucleoid is degraded, the complex is free to diffuse and we expect the principal direction to be the major axis of the cell. The PCA then gets rid of boundary effects of the shortest direction of the cell, and thus offers a more stable MSD for longer lag times. However, under conditions where the nucleoid is not degraded, we cannot extract the cell orientation from the PCA due to the confined movement of the complexes imposed by ParA (see figure below), but we still used the PCA so that all MSDs are 1D and direct comparisons were easier.

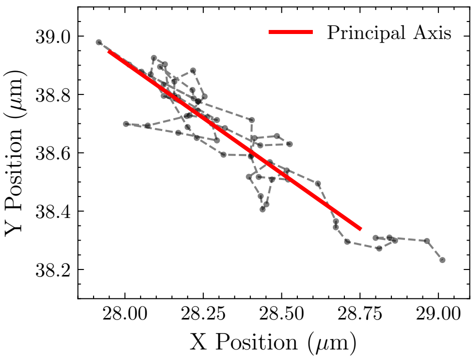

(a)

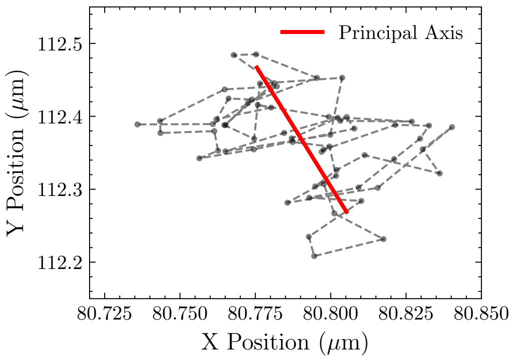

(b)

**Figure :** Two typical trajectories of complexes with (a) and without (b) degradation of the nucleoid. The principal direction obtained from PCA is represented by the red line.

The MSD is averaged over $n$ independent tracks and then fitted at short lag time ($\Delta t \cdot \delta t < t_{cutoff}$) with a power law:

$$\mathrm{MSD}(\Delta t) = D_\alpha \cdot t^\alpha \tag{5}$$

The parameters are summarized in the following table :

| | MSD_F_wo_chrom | MSD_F+chrom | MSD_mF_wo_chrom | MSD_mF+chrom |
|---|---|---|---|---|
| n | 230 | 707 | 274 | 427 |
| $t_{cutoff}(s)$ | 2.5 | 30 | 2.5 | 30 |
| $\alpha$ | $0.92 \pm 0.02$ | $0.807 \pm 0.003$ | $0.95 \pm 0.03$ | $0.880 \pm 0.005$ |
| $D_\alpha \left(\mu m^2 \cdot s^{-\alpha}\right)$ | $(19.1 \pm 0.4) \cdot 10^{-3}$ | $(19.2 \pm 0.1) \cdot 10^{-4}$ | $(43 \pm 1) \cdot 10^{-3}$ | $(11.3 \pm 0.2) \cdot 10^{-4}$ |

### Polymer model and coarse-graining

As described experimentally (Guilhas et al., 2020), ParB-DNA complexes have approximately a spherical shape of $37 \pm 5nm$. Each plasmid is represented by either two linear self-avoiding chains (SAWs) of $N$ monomers or by a single star polymer with $k = 4$ arms of length $L = N/2$, with a central junction. All monomers correspond to a DNA length of $a = 50\,\text{nm}$ according to the persistence length of DNA. The linear configuration represents the initial separated state where DNA from each condensate behaves as independent polymer chains. The star polymer configuration models the fused state where DNA from both condensates is constrained within a single droplet, with the central junction representing the geometric constraint imposed by fusion. This comparison provides an estimate of the entropy penalty upon fusion.

The partition functions to compute the entropic term are :

$$Z_{\text{linear}} = \left(\mu_{\text{SAW}}\, N^{\gamma_{\text{SAW}}-1}\right)^2$$

And for stars polymers (Duplantier, 1986) :

$$Z_{\text{star}} = \mu_{\text{SAW}}^{kL}\, L^{\sigma_k + k\,\sigma_1}$$

With their approximate values in $d = 3$,

$$\mu_{\text{SAW}} = 2.638159, \quad \gamma_{\text{SAW}} = 1.157, \quad \sigma_4 = -0.48, \quad \sigma_1 = 0.08$$

### Free-energy balance $\Delta F$

We define

$$\Delta F \;=\; F_{\text{fused}} - F_{\text{init}} \;=\; \left[\gamma\, A_{\text{fused}} - k_B T \ln Z_{\text{star}}\right] \;-\; \left[2\gamma\, A_1 - k_B T \ln Z_{\text{linear}}\right].$$

The areas are

$$A_1 = 4\pi R^2, \quad A_{\text{fused}} = 4\pi\,(2R^3)^{2/3}, \quad \Delta A = 2A_1 - A_{\text{fused}},$$

and the entropy difference is

$$\Delta S_{\text{plasmid}} = k_B\left(\ln Z_{\text{star}} - \ln Z_{\text{linear}}\right).$$

At the fusion–separation transition ($\Delta F = 0$), the critical surface tension is given by

$$\gamma_{\text{crit}} = \frac{k_B T\, \Delta S_{\text{plasmid}}}{\Delta A}.$$

One model-unit of surface tension corresponds to

$$\frac{k_B T}{a^2} = \frac{4.1 \times 10^{-21}\,\text{J}}{(50 \times 10^{-9}\,\text{m})^2} \approx 1.64 \times 10^{-6}\,\text{N/m} = 1.64\,\mu\text{N/m}.$$

Thus

$$\gamma_{\text{phys}} = \gamma_{\text{model}} \times 1.64\,\mu\text{N/m}.$$

Error-bar propagation are :

$$\frac{\sigma_{\Delta A}}{\Delta A} \approx \left|\frac{d\Delta A}{dR}\right| \frac{\sigma_R}{\Delta A} = 2\,\frac{\sigma_R}{R} \approx 0.267.$$

Since $\gamma_{\text{crit}} \propto 1/\Delta A$,

$$\frac{\sigma_\gamma}{\gamma} = \frac{\sigma_{\Delta A}}{\Delta A}, \quad \sigma_{\gamma_{\text{phys}}} = \gamma_{\text{phys}}\,\frac{\sigma_{\Delta A}}{\Delta A}.$$
